# FSP1 and histone deacetylases suppress cancer persister cell ferroptosis

**DOI:** 10.1101/2025.08.21.671520

**Authors:** Masayoshi Higuchi, August F. Williams, Anna E. Stuhlfire, Ariel H. Nguyen, David A. G. Gervasio, Claire E. Turkal, Suejean Chon, Matthew J. Hangauer

**Author notes:** Correspondence to (M.J.H.). These authors contributed equally to this work (Masayoshi Higuchi, August F. Williams, Anna E. Stuhlfire).

## Abstract

Cancer persister cells populate minimal residual disease and contribute to acquired drug resistance. We previously discovered that persister cells are sensitized to ferroptosis. However, our understanding of this emergent persister cell vulnerability remains limited, impeding ferroptosis drug development efforts. Here, we sought to understand key factors which govern persister cell ferroptosis to inform combinatorial treatment strategies. We found that persister cells can downregulate oxidative phosphorylation, a key source of reactive oxygen species, to avoid death from GPX4 inhibition. However, this can be overcome by pretreatment with clinically available histone deacetylase inhibitors which induce reactive oxygen species in persister cells and synergize with GPX4 inhibition. Furthermore, we found that while levels of iron, glutathione, and antioxidant genes are not universally dysregulated in persister cells, persister cells consistently downregulate alternative ferroptosis suppressor FSP1 and rely upon residual FSP1 to survive GPX4 inhibition. These findings reveal new strategies to eliminate persister cells by combining GPX4 inhibitors with histone deacetylase or FSP1 inhibitors.

## Introduction

Drug-tolerant cancer persister cells survive cytotoxic drug treatments and populate minimal residual disease, undergo mutagenesis, and may contribute to acquired resistance and tumor recurrence (*1*). Therapeutic eradication of persister cells may increase the durability of responses to existing cancer treatments. Unfortunately, there are no clinically approved therapies which are intended to target persister cells. Identification of a robust persister cell drug target has been challenging because the majority of known persister cell vulnerabilities are restricted to specific cell types and drug treatments (*2–11*). One notable exception is ferroptosis. We previously discovered that persister cells from multiple tumor types and treatments are vulnerable to ferroptosis which can be induced with inhibitors of the lipid hydroperoxidase GPX4 (*3*). There are multiple drug development efforts underway targeting GPX4, but GPX4 is an essential enzyme raising toxicity concerns and thus far there have been no reported GPX4 inhibitors with potent in vivo efficacy. Also, a GPX4 inhibitor must not only achieve bioavailability, it may also need enhanced potency because ferroptosis sensitivity in vivo is diminished compared to cell culture (*12–14*).

Alternative approaches to induce ferroptosis in persister cells may overcome these challenges. Unfortunately, it remains poorly understood why persister cells are selectively sensitized to ferroptosis despite a variety of factors which promote ferroptosis sensitivity being known in other contexts (*15*). It was recently postulated that persister cells utilize iron to survive, similar to cancer stem cells and cells undergoing epithelial to mesenchymal transition (*4, 16–18*), and that this results in enhanced sensitivity to ferroptosis (*19*). However, we previously found that persister cells have decreased rather than increased total cellular iron levels,(*3*) though iron has been reported to be increased in a lung cancer persister model (*20*) and it remains possible that subcellular labile iron pools, such as within the lysosome,(*21, 22*) differ in persister cells. Another possibility is that persister cells are deficient in antioxidant defenses. Indeed, we previously found that persister cells have lower basal glutathione (GSH) levels, which was also observed in another persister cell model,(*20*) and that GSH is rapidly further depleted and reactive oxygen species (ROS) dramatically increased upon GPX4 inhibition in persister but not drug naïve parental cells (*3*). However, there is relatively modest rescue from persister cell ferroptosis by GSH replenishment indicating low GSH may not be necessary for ferroptosis sensitivity (*3*). Taken together, these observations show that while cellular iron and GSH levels influence ferroptosis, neither fully explain why persister cells are sensitized to ferroptosis.

In this study, we sought to explore alternative explanations for persister cell ferroptosis sensitivity. Persister cells have been previously reported to preferentially depend on oxidative phosphorylation, a major source of ROS, instead of glycolysis metabolism (*23–25*). Here, we found that persister cell dependence on oxidative phosphorylation is an Achilles heel because it generates oxidative stress which sensitizes to ferroptosis. This vulnerability can be exacerbated by treatment of persister cells with otherwise nontoxic concentrations of clinically available histone deacetylase (HDAC) inhibitors which increase oxidative phosphorylation and ROS and further sensitize persister cells, but not parental cells, to ferroptosis. Also, while persister cells are not universally sensitive to FSP1 inhibition despite decreased FSP1 expression, persister cells which survive ferroptosis induced by GPX4 inhibition become sensitized to inhibition of FSP1. Therefore, HDAC inhibitors or FSP1 inhibitors may be combined with GPX4 inhibitors to selectively enhance persister cell ferroptosis. These findings provide insight into why cancer persister cells are sensitized to ferroptosis and reveal potential combinatorial treatment strategies to enhance persister cell ferroptosis while potentially sparing other cells.

## Results

### Oxidative phosphorylation contributes to persister cell sensitivity to ferroptosis

We reasoned that analysis of the subpopulation of persister cells which survive partially lethal GPX4 inhibition may reveal why persister cells are sensitized to ferroptosis. We therefore performed single cell RNA sequencing (scRNAseq) on EGFR mutant PC9 non-small cell lung cancer persister cells, derived from 10 days of treatment with EGFR inhibitor erlotinib, which were subsequently treated with GPX4 inhibitor RSL3 for 24 hours resulting in partial death and effectively purifying surviving persister cells. RSL3-treated persister cells which survived RSL3 treatment were nearly completely separated from persister cells which were not treated with RSL3 on a UMAP plot, indicating surviving cells were altered by RSL3 exposure (Fig. 1A). In contrast, drug naïve PC9 parental cells treated with the same concentration and duration of RSL3, which is nontoxic to parental cells, entirely overlapped untreated parental cells indicating that GPX4 inhibition has minimal effect on parental cells (Fig. 1A). These observations are consistent with our prior findings that persister cells are sensitized to ferroptosis relative to parental cells (*3*).

**Fig. 1.**
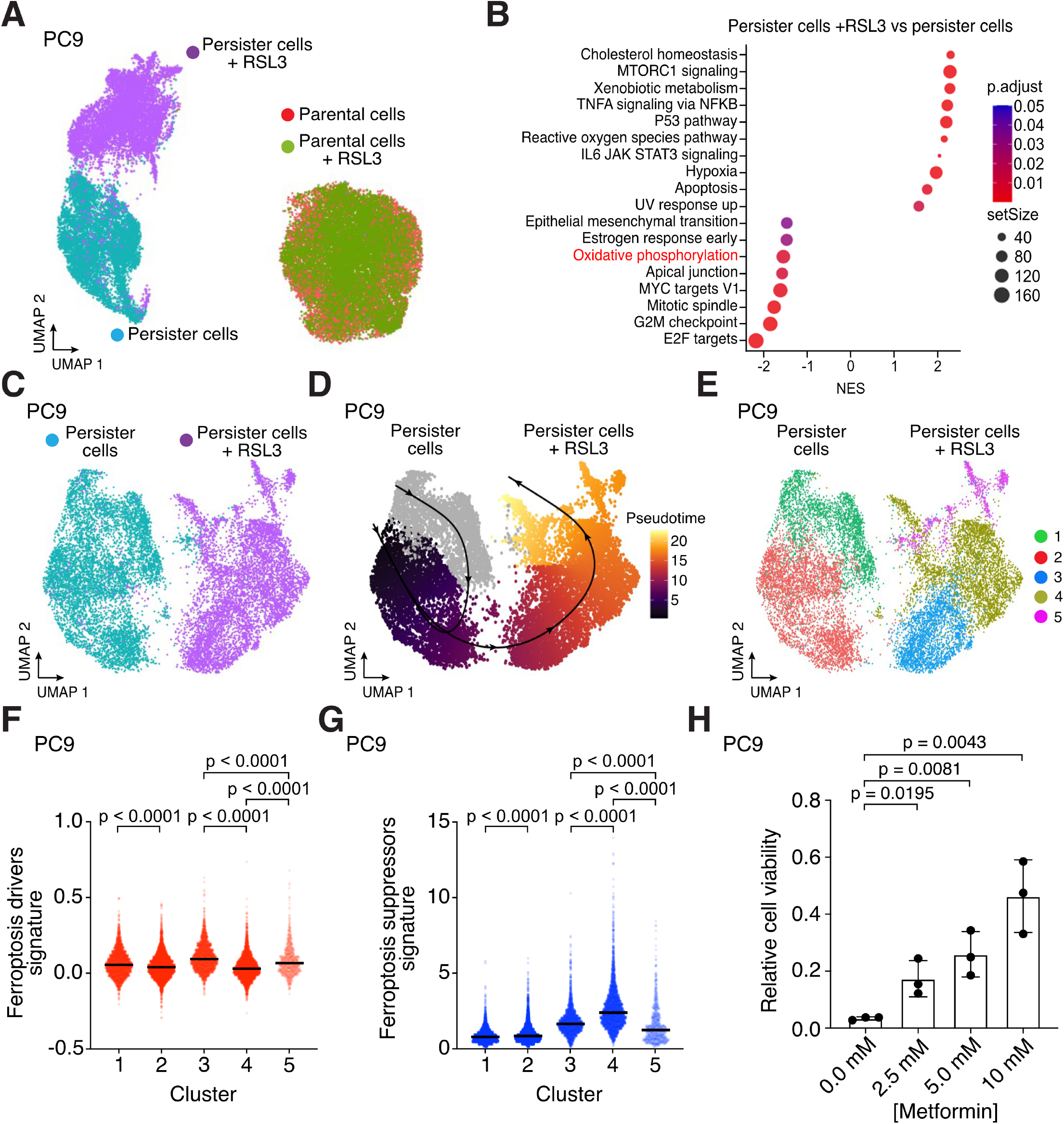
Oxidative phosphorylation contributes to ferroptosis sensitization in cancer persister cells. (**A**) UMAP of PC9 parental and persister cells treated with or without 1 μM RSL3 for 24 hours. (**B**) Enriched Hallmarks gene sets in persister cells treated with and without RSL3. Positive values indicate enrichment in persister cells treated with RSL3. (**C**) UMAP of PC9 persister cells treated with and without RSL3. (**D**) Pseudotime analysis of PC9 persister cells treated with and without RSL3. Solid black line indicates the estimated transitional trajectory across cell states. (**E**) UMAP of PC9 persister cells treated with and without RSL3 colored by cluster. (**F**) Ferroptosis-drivers gene set signature score across clusters in (**E**). (**G**) Ferroptosis-suppressors gene set signature score across clusters in (**E**). (**F** and **G**) P values calculated with Mann-Whitney test. (**H**) PC9 persister cells derived from erlotinib plus metformin and then treated with 500 nM RSL3 for 24 hours. *n* = 3 biological replicates; mean ± s.d. is shown; P values calculated with two-tailed Student’s t-test.

Gene set enrichment analysis of persister cells revealed pathways known to protect from ferroptosis were enriched among persister cells which survived RSL3 treatment including cholesterol homeostasis (*26*), MTORC signaling (*27*), xenobiotic (glutathione) metabolism (*28*), TNF*α* signaling via NFκB (*29*), and p53 signaling (*30*) (Fig. 1B, and tables S1 and S2). The epithelial-to-mesenchymal (EMT) gene set was also downregulated in cells surviving RSL3 treatment, consistent with prior reports that mesenchymal cancer cells are sensitized to ferroptosis (Fig. 1B) (*4*). Furthermore, a YAP gene set signature was depleted among persister cells surviving RSL3 treatment (fig. S1 and table S2). It was previously reported that loss of YAP signaling upon cell contact-induced activation of the Hippo pathway protects cells from ferroptosis (*13*). This raised the possibility that low cell density may explain persister cell sensitivity to ferroptosis. To test this, we generated “high density” PC9 persister cells and BRAF and MEK inhibitor-derived BRAF V600E A375 melanoma persister cells by treating with lower drug concentrations for 10 days resulting in fully confluent persister cells and found that high density persister cells remain selectively sensitized to ferroptosis versus density-matched parental cells of equal or lower confluency (fig. S2). Therefore, persister cell sensitization to ferroptosis is not merely due to low cell density.

Pseudotime analysis of the scRNAseq data revealed a trajectory from persister cells which were not treated with RSL3 toward cells which survived RSL3 treatment (Fig. 1, C to E). Persister cells which survived RSL3 treatment separated into three clusters. Along the pseudotime trajectory, the first RSL3-treated cluster (Cluster 3) is characterized by relatively high expression of a ferroptosis-driver gene set signature and lower ferroptosis-suppressor signature compared to the other RSL3-treated clusters (*31*), suggesting that this cluster contains relatively ferroptosis-sensitive cells (Fig. 1, D to G). In contrast, the next RSL3-treated persister cell cluster along the trajectory (Cluster 4) has lower expression of the ferroptosis-driver gene set and higher expression of the ferroptosis-suppressor gene set suggesting that this population is more protected from ferroptosis (Fig. 1, D to G). The third RSL3-treated cluster (Cluster 5) has relatively few cells and exhibited a high ferroptosis-driver signature and low ferroptosis-suppressor signature suggesting this is also a ferroptosis sensitive population (Fig. 1, E to G and Tables S3 and S4).

Among the gene sets differentially expressed in ferroptosis-resistant Cluster 4 versus the other two RSL3-treated clusters, we observed decreased expression of the oxidative phosphorylation gene set (Tables S3 and S4). Given that the bulk population of persister cells treated with RSL3 also show decreased expression of the oxidative phosphorylation gene set (Fig. 1B), we hypothesized that oxidative phosphorylation, a persister cell dependency and ROS source (*23, 24*), contributes to ferroptosis sensitization of persister cells. To test this hypothesis, we utilized metformin, an inhibitor of oxidative phosphorylation which blocks mitochondrial electron transport chain complex I, and found that while metformin is partially toxic to persister cells (fig. S3), consistent with oxidative phosphorylation as an important persister cell energy source (*23–25*), the surviving persister cells displayed metformin dose-dependent protection against RSL3 indicating inhibition of oxidative phosphorylation protects from ferroptosis (Fig. 1H). Together, these observations show that oxidative phosphorylation contributes to persister cell ferroptosis sensitivity.

### FSP1 expression determines persister cell GPX4 sensitivity

We previously found that HER2 amplified BT474 breast cancer persister cells which are sensitized to ferroptosis have decreased expression of NRF2 target genes reflecting a disabled antioxidant state (*3*). Here, we found that the PC9 persister cells which survive RSL3 are enriched for antioxidant genes in the reactive oxygen species pathway (Fig. 1B and table S2). Furthermore, we found that PC9 persister cells which survive RSL3 treatment are enriched for the NRF2 gene set consistent with NRF2 acting as a master antioxidant transcription factor in persister cells (fig. S1). We also found that NRF2 protein levels are decreased in all tested persister cell types relative to parental cells (Fig. 2A). However, the role NRF2 plays in ferroptosis protection is controversial (*15*) and, unlike BT474 persister cells, we found that PC9 persister cells show enrichment rather than depletion of the NRF2 gene set compared to parental cells (fig. S4, tables S5 and S6). Also, the NRF2 negative regulator KEAP1 is consistently decreased rather than increased across persister cell types reflecting potentially complex regulation (Fig. 2A). Therefore, while NRF2 activity may protect from ferroptosis, loss of NRF2 activity does not appear to universally contribute to ferroptosis sensitivity in all persister cell types.

**Fig. 2.**
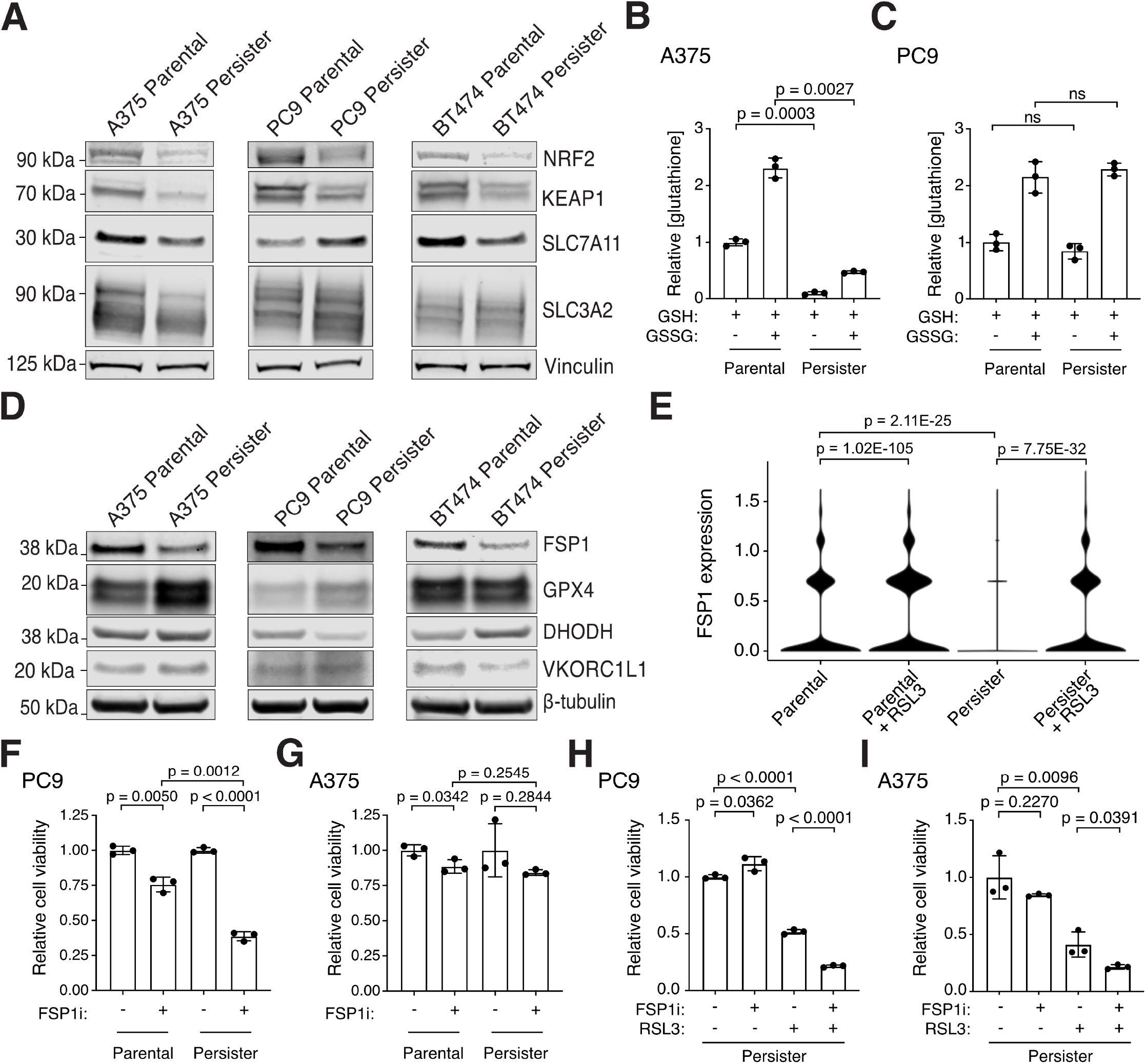
Persister cells have variable antioxidant deficiencies and depend on FSP1 to survive GPX4 inhibition. (**A**) Parental and persister cells analyzed for NRF2, KEAP1, and system xc^−^ components SLC7A11 and SLC3A2 expression. (**B** and **C**) A375 and PC9 parental and persister cells analyzed for reduced GSH or total GSH (GSSG) levels. (**D**) Protein expression of ferroptosis suppressor genes in parental and persister cells. (**E**) FSP1 mRNA expression in PC9 parental and persister cells treated with and without RSL3. P values calculated with the Wilcoxon Rank Sum test with Bonferroni correction. (**F** and **G**) PC9 and A375 parental and persister cells treated with and without 5 μM FSP1 inhibitor. (**H** and **I**) PC9 and A375 persister cells co-treated with 1 μM FSP1 inhibitor, 50 nM RSL3, or both for 24 hours. (**B, C, F** to **I**) *n* = 3 biological replicates; mean ± s.d. is shown; P values calculated with two-tailed Student’s t-test.

We previously found that BT474 persister cells exhibit decreased expression of SLC7A11, a component of system x_C_^-^, which is responsible for the transport of the GSH precursor cystine into the cell (*15*), and have decreased GSH levels (*3*). We also found here that PC9 persister cells with increased xenobiotic metabolism, which is related to GSH function, were selected for among persister cells which survived RSL3 treatment (Fig. 1B). Together with our prior observation that GSH is rapidly depleted in BT474 persister but not parental cells upon GPX4 inhibition, and that N-acetyl cysteine treatment partially rescues from ferroptosis (*3*), these data support that GSH counteracts ferroptosis in persister cells. However, relative levels of system x_C_^-^ components SLC7A11 and SLC3A2 proteins were variable across persister cell types and while A375 persister cells have reduced GSH levels compared to parental cells, similar to BT474 persister cells, PC9 persister cells do not (Fig. 2, A to C). Therefore, while GSH protects from ferroptosis, decreased persister cell GSH is not a general cause of cancer persister cell ferroptosis sensitivity across all persister cell types.

We next investigated the protein levels of select ferroptosis suppressing enzymes including GPX4 (*32*), FSP1 (*33, 34*), DHODH (*35*) and VKORC1L1 (*36*) to determine if loss of any of these factors could explain the persister cell dependence on GPX4. Interestingly, FSP1 protein expression was decreased across all persister cell models, while GPX4, DHODH and VKORC1L1 levels were variable (Fig. 2D). Furthermore, while FSP1 mRNA expression was decreased in PC9 persister cells compared to parental cells, it increased in persister cells which survive GPX4 inhibition (Fig. 2E). Based on this, we hypothesized that persister cells which survive GPX4 inhibition may become reliant upon FSP1 to survive. We therefore tested whether GPX4 inhibitor-treated persister cells are sensitized to FSP1 inhibition. While FSP1 inhibitor alone had variable effects on persister cells (Fig. 2, F and G), we found that combining FSP1 inhibitor with GPX4 inhibitor increased persister cell death significantly across persister cell models including in A375 persister cells in which FSP1 inhibitor alone is nontoxic (Fig. 2, H and I). Therefore, persister cells depend on FSP1 when GPX4 is inhibited. These observations support combining inhibition of ferroptosis suppressor FSP1 with GPX4 inhibition to enhance persister cell ferroptosis. We next focused on whether a pro-ferroptosis factor may be increased as an alternative approach to target persister cells for ferroptosis.

### HDAC inhibitors synergize with GPX4 inhibition to induce persister cell death

Histone deacetylase (HDAC) inhibitors were recently shown to enhance ferroptosis in combination with inhibitors of system x_C_^-^ or GPX4 in other contexts (*37–44*). We therefore explored whether HDAC inhibition also promotes persister cell ferroptosis. Remarkably, we found that clinically utilized pan-HDAC inhibitors panobinostat and vorinostat synergize with GPX4 inhibitors to induce cell death in all tested lung, melanoma and breast cancer persister cell models but not in parental cells (Fig. 3, A to F and fig. S5A). We also found synergy between RSL3 and another epigenetic modifier, BRD4 inhibitor JQ1 (fig. S5B). Furthermore, pretreatment with a nontoxic concentration of HDAC inhibitor sensitized persister cells to subsequent GPX4 inhibitor treatment (Fig. 3, G to J and fig. S5, C to F). Therefore, modulation of persister cell epigenetic states can sensitize to ferroptosis.

**Fig. 3.**
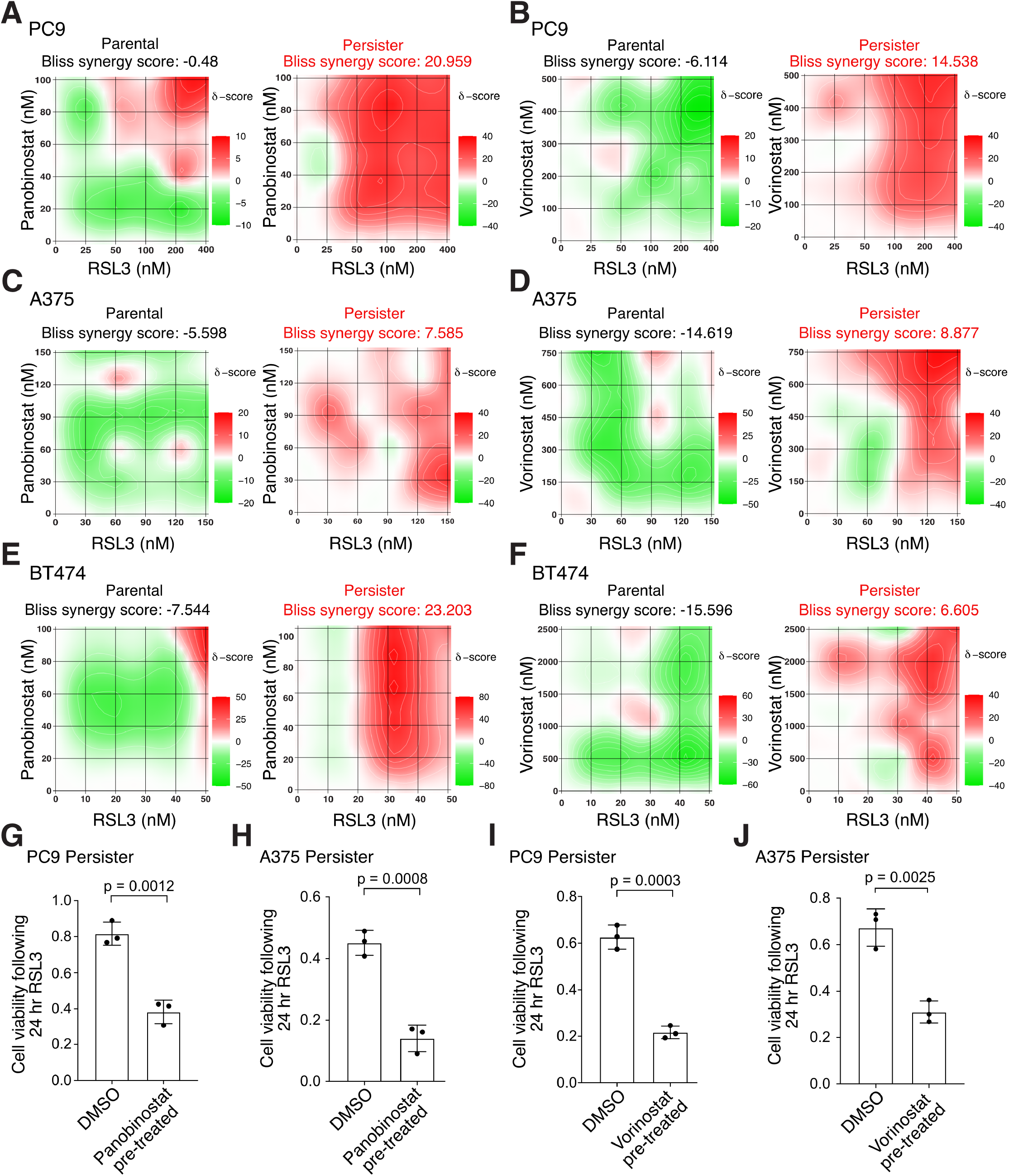
HDAC inhibition synergizes with GPX4 inhibition to selectively kill persister cells. (**A** to **F**) Heatmaps of synergy between GPX4 inhibitor RSL3 and HDAC inhibitors panobinostat and vorinostat following 24 hour cotreatment. Bliss synergy score calculated with SynergyFinder 3.0. Red color and positive scores indicate synergy, green color and negative scores indicate buffering. (**A** and **B**) PC9 parental cells and persister cells derived from 50 nM erlotinib. (**C** and **D**) A375 parental cells and persister cells derived from 10 nM dabrafenib with 1 nM trametinib. (**E** and **F**) BT474 parental cells and persister cells derived from 2 μM lapatinib. (**G** to **J**) Pre-derived PC9 or A375 persister cells were treated for 48 hours with a nontoxic concentration of HDAC inhibitor (see fig. S5), rinsed, and then treated with RSL3 for 24 hours while maintained under targeted therapy treatment. Data normalized to untreated persister cells. Concentration and HDAC inhibitor used: (**G**) 7.5 nM panobinostat, (**H**) 5 nM panobinostat, (**I**) 100 nM vorinostat, (**J**) 1 μM vorinostat. RSL3 concentrations used: (**G**) 150 nM, (**H**) 100 nM, (**I**) 150 nM, (**J**) 80 nM. *n* = 3 biological replicates; mean ± s.d. is shown; P values calculated with two-tailed Student’s t-test.

We performed scRNAseq on panobinostat-treated PC9 persister cells and found that, unlike RSL3 treatment which solely affects persister cell transcriptomes (Fig. 1A), panobinostat causes broad transcriptional changes to both parental and persister cells (Fig. 4A). Xenobiotic metabolism was an enriched gene set upon HDAC inhibitor treatment of persister cells (Fig. 4B). We therefore tested whether HDAC inhibitor treatment affects GSH levels in persister cells. However, HDAC inhibitor did not affect persister cell GSH levels (Fig. 4, C and D) and addition of excess GSH did not rescue persister cells from HDAC inhibitor-induced ferroptosis sensitization (Fig. 4E). This observation is consistent with our earlier data showing that GSH replenishment is not sufficient to rescue persister cells from ferroptosis (*3*). Furthermore, while the heme metabolism gene set was enriched in persister cells upon HDAC inhibition indicating a potential role for iron (Fig. 4B), we found that HDAC inhibition lowered rather than increased the labile iron content in persister cells (Fig. 4F). Therefore, HDAC inhibitor treatment enhances persister cell sensitization to ferroptosis without altering GSH or iron levels.

**Fig. 4.**
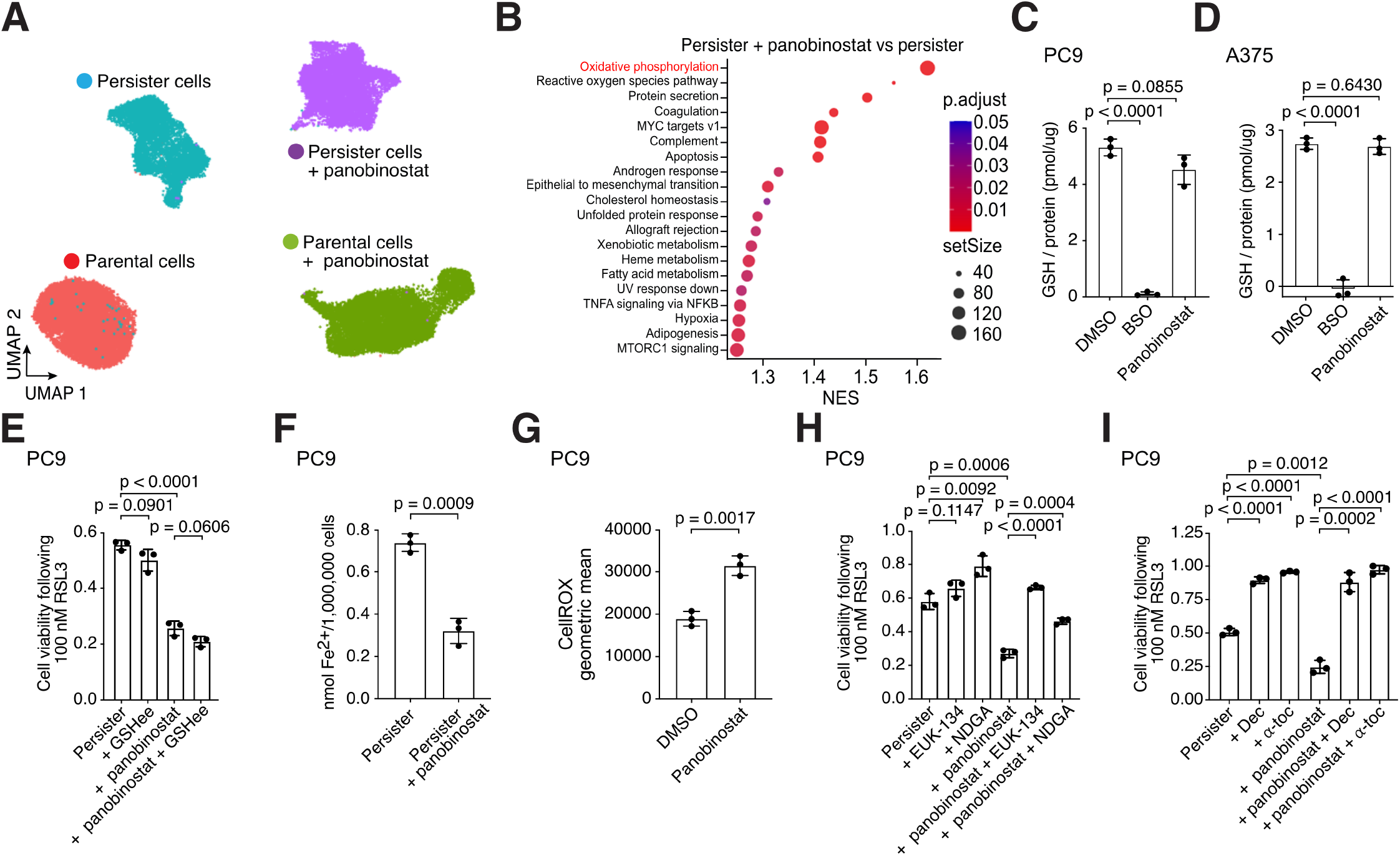
HDAC inhibitors induce persister cell oxidative stress to sensitize persister cells to ferroptosis. (**A**) UMAP of PC9 parental and persister cells treated with or without HDAC inhibitor panobinostat for 48 hours. (**B**) Treatment of PC9 persister cells with panobinostat enriches for oxidative phosphorylation genes. (**C** and **D**) Treatment with panobinostat does not decrease GSH levels in PC9 or A375 persister cells. 1 mM buthionine sulfoximine (BSO) was used as a positive control for GSH depletion. (**E**) Ferroptosis sensitization of PC9 persister cells from treatment with panobinostat is not inhibited by glutathione ethyl ester (GSHee, 1 mM). (**F**) PC9 persister cell treatment with panobinostat decreases rather than increases intracellular iron. (**G**) Panobinostat treatment of PC9 persister cells increases ROS. (**H** and **I**) PC9 persister cells treated with panobinostat are rescued from RSL3 treatment with the antioxidants EUK-134 (10 μM), nordihydroguaiaretic acid (NDGA, 5 μM), decylubiquinone (Dec, 7.5 μM), and *α*-tocopherol (*α*-toc, 1.5 mM). Panobinostat concentrations used in PC9 cells: 7.5 nM (**A** to **C** and **F** to **G**) or 2.5 nM (**E** and **H** to **I**); in A375 cells: 5 nM (**D**). (**C** to **I**) *n* = 3 biological replicates; mean ± s.d. is shown; P values calculated with two-tailed Student’s t-test.

Instead, consistent with our earlier observations that oxidative phosphorylation underlies persister cell ferroptosis sensitivity, we found that oxidative phosphorylation and ROS were the two top Hallmarks gene sets enriched upon HDAC inhibitor treatment in persister cells (Fig. 4B, and tables S7 and S8) and that HDAC inhibition increases persister cell ROS (Fig. 4G). Furthermore, diverse soluble and lipophilic antioxidants including EUK-134, nordihydroguaiaretic acid (NDGA), decylubiquinone and *α*-tocopherol negated HDAC inhibitor-induced ferroptosis sensitization (Fig. 4, H and I). Therefore, HDAC inhibitor treatment induces oxidative stress in persister cells which sensitizes to ferroptosis.

## Discussion

We previously reported that cancer persister cells are sensitized to ferroptosis. Since that time, accumulating data have reinforced this finding (*1, 20, 22, 45–49*), yet there has been minimal progress in understanding why ferroptosis is an emergent persister cell vulnerability. Furthermore, there has yet to be developed a potent chemical inducer of ferroptosis with strong efficacy and minimal toxicity in vivo. An improved mechanistic understanding of the persister cell ferroptosis vulnerability may help identify new therapeutic approaches which overcome current hurdles. We reasoned that transcriptomic changes within persister cells which survive brief ferroptotic stress may implicate key pathways which govern persister cell ferroptosis sensitivity. Consistent with ferroptosis as a persister cell-selective vulnerability, we found that while persister cells which survive GPX4 inhibitor exposure are transcriptionally distinct from other persister cells, drug naïve parental cells are transcriptionally unaffected by GPX4 inhibition. Furthermore, we found that oxidative phosphorylation genes are depleted among persister cells which survive RSL3 treatment indicating that oxidative phosphorylation may contribute to persister cell ferroptotic death. Indeed, persister cells with chemically inhibited oxidative phosphorylation, via metformin treatment, are partially protected from ferroptosis. Therefore, oxidative phosphorylation, a known persister cell dependency and major source of cellular ROS (*1, 23, 24, 50*), is a driver of persister cell ferroptosis sensitivity.

While increased ROS production due to a dependence on oxidative phosphorylation can promote ferroptosis, a disabled antioxidant program may also sensitize persister cells to ferroptosis. We previously found that HER2 amplified BT474 breast cancer persister cells have a broadly disabled antioxidant program with diminished expression of NRF2 target genes including system x_C_^-^ components and decreased levels of GSH and NADPH(*3*). Interestingly, while each of the lung, melanoma and breast cancer persister cell types we analyzed here are deficient in one or more anti-ferroptotic factors including NRF2, GSH or in anti-ferroptotic enzymes GPX4, DHODH or VKORC1L1, all were deficient in FSP1. Therefore, while persister cells have heterogeneous and cell type-specific antioxidant deficiencies, decreased FSP1 expression may underly the persister cell dependence on GPX4 for protection from ferroptosis. Furthermore, we found that multiple persister cell types depend on FSP1 to survive in the presence of GPX4 inhibitor treatment, indicating that FSP1 is a crucial factor for persister cells to survive ferroptotic stress.

We also searched for a new strategy to enhance persister cell ferroptosis. We found that the FDA-approved pan-HDAC inhibitors panobinostat and vorinostat synergize with GPX4 inhibitors to enhance persister cell ferroptosis. Although their clinical efficacy has been limited when used as monotherapies or showed mixed results when used in combination in solid tumors (*51*), our data suggest that if applied to minimal residual disease, HDAC inhibitors may prime persister cells for ferroptosis. Interestingly, HDAC inhibitor treatment alone has been shown to increase lipid peroxidation and modulate ferroptosis-related genes in gastric cancer cell lines (*52*) and studies in acute myeloid leukemia and colorectal cancer models found a synergistic effect combining HDAC inhibitors with GPX4 inhibitors (*37, 38*). Others have also found increased ferroptosis sensitization when combining HDAC inhibitors with the system x_C_^-^ inhibitor erastin (*39–44*). However, while increased iron uptake (*37, 44*) and decreased GSH levels from a reduction in system x_C_^-^ factors (*43, 52*) were suggested as the HDAC inhibitor-induced ferroptosis sensitization mechanisms, we found that HDAC inhibitor decreased the iron content and did not impact GSH levels in cancer persister cells. Instead, in persister cells, we found that HDAC inhibitors upregulate oxidative phosphorylation and increase ROS levels. As HDAC inhibitors have broad effects on gene expression, further study will be needed to determine precisely how HDAC inhibitors increase persister cell oxidative stress including determining whether this is due to a nonspecific stress response or is facilitated by specific derepressed genes. Regardless of how HDAC inhibitors promote persister cell oxidative stress, our finding that nontoxic concentrations of clinically available HDAC inhibitors specifically sensitize persister cells to ferroptosis highlights a potential combinatorial treatment approach with HDAC inhibitor and a bioavailable and potent ferroptosis inducer, once developed, to eradicate persister cells and prevent acquired resistance. Furthermore, hybrid molecules that target both HDACs and induce ferroptosis simultaneously may confer unique responses that warrant additional testing (*53*).

While a full understanding of persister cell sensitivity to ferroptosis will require further studies of their features such as phospholipid composition (*17*), membrane rupture (*54*) and repair (*55, 56*), here we have illuminated a key feature of persister cell ferroptosis sensitivity based on their dependence on oxidative phosphorylation which contributes to elevated persister cell oxidative stress (Fig. 5). Furthermore, we found that this vulnerability can be exacerbated by treatment with HDAC inhibitors and that persister cells which survive GPX4 inhibition can be further targeted with FSP1 inhibition. Given the concern that GPX4 inhibitors may have unacceptable toxicity in humans (*57–59*), discovery of combinatorial approaches such as these to selectively sensitize cancer cell populations may be critical to enable the realization of a clinically effective and safe ferroptosis-based drug treatment for cancer.

**Fig. 5.**
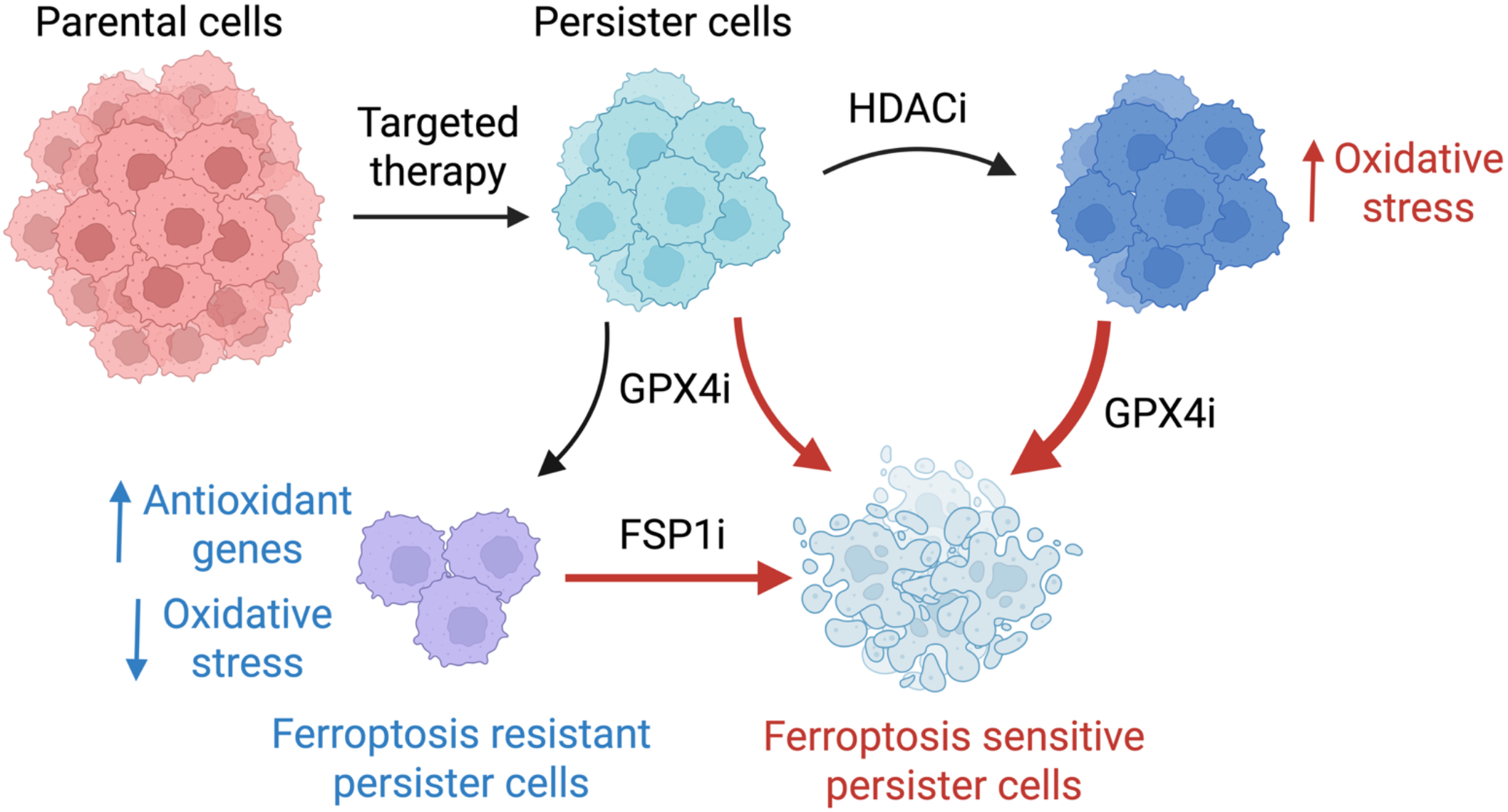
Enhancing persister cell ferroptosis with FSP1 and HDAC inhibition. Cancer persister cells can decrease oxidative stress to survive GPX4 inhibition (GPX4i). However, GPX4i-tolerant persister cells become dependent on the alternative ferroptosis suppressor enzyme FSP1 to survive and addition of FSP1 inhibitor (FSP1i) increases persister cell ferroptotic death. Furthermore, persister cell oxidative stress is increased by nontoxic pre- or co-treatment with clinically available HDAC inhibitors resulting in synergistic persister cell ferroptosis in combination with GPX4 inhibitor (GPX4i). Our findings reveal new approaches to selectively enhance persister cell ferroptosis while potentially sparing other cells.

## Materials and Methods

### Cell culture

PC9 cells were provided by the Altschuler and Wu lab at University of California San Francisco. BT474 (HTB-20) and A375 (CRL-1619) cells were purchased from ATCC. PC9 cells were cultured in RPMI-1640 (Gibco, 11875093) supplemented with 5% fetal bovine serum (FBS) and 1% Antimycotic/Antibiotic (AA) (ThermoFisher Scientific). A375 cells were cultured in DMEM (high glucose, Gibco, 11965092) supplemented with 10% FBS and 1% AA. BT474 cells were cultured in RPMI-1640 supplemented with 10% FBS and 1% AA. All cells were incubated at 5% CO2 and 37 °C. Cell lines were split with 0.25% trypsin-EDTA (ThermoFisher Scientific, 25200056). Cell line identities were confirmed with STR profiling at the UC Berkeley Cell Culture Facility. All cell lines regularly tested negative for mycoplasma throughout these investigations using the Lonza Mycoalert Mycoplasma Detection Kit (Lonza, LT07-318).

### Chemicals

ML210 (SML0521), (1S,3R)-RSL3 (SML2234), decylubiquinone (D7911), L-buthionine-sulfoximine (BSO) (B2515), glutathione reduced ethyl ester (CH6H9A56C7FA), (±)-α-tocopherol (258024), and EUK-134 (SML0743) were purchased from Sigma-Aldrich. Lapatinib (S2111), erlotinib (S7786), and iFSP1 (S9663) were purchased from Selleck Chemicals. JQ1 (11187) and panobinostat (13280) were purchased from Cayman Chemical. Vorinostat (HY-10221), dabrafenib (HY-14660), trametinib (HY-10999), and metformin (HY-B0627) were purchased from MedChemExpress. All chemicals except for BSO and metformin, which were dissolved and stored in media, were stored as stock solutions in DMSO (Life Technologies) or cell culture grade water (Corning).

### Persister cell derivation

High density PC9 and A375 persister cells were used for all figures except Fig. 1H and western blots where low density persister cells were used. BT474 low density persister cells were used for all experiments. High and low density persister cells, and density matched parental cells, were derived for each cell line as follows. Low density PC9 parental cells were seeded at 5,200/well in 96 well plates and high density PC9 parental cells were seeded at 21,000/well in 96 well plates. Low density A375 parental cells were seeded at 4,000/well in 96 well plates and high density A375 parental cells were seeded at 8,000/well in 96 well plates. For PC9 and A375, experiments were performed 24 hours after seeding. Low density BT474 parental cells were seeded at 3,000/well 96 well plates and experiments were performed 72 hours after seeding. To derive PC9 persister cells at low cell density, cells were seeded at 1,800/well in 96 well plates and after 24 hours were subjected to continuous treatment with 2.5 μM erlotinib (Selleck chemicals) for 10 days. To derive PC9 persister cells at high cell density, cells were seeded at 11,000/well in 96 well plates and after 24 hours were subjected to continuous treatment with 70 nM erlotinib for 10 days unless otherwise stated. To derive A375 persister cells at low cell density, cells were seeded at 2,300/well in 96 well plates and after 24 hours were subjected to continuous treatment with 500 nM dabrafenib (Selleck chemicals) and 50 nM trametinib (Selleck chemicals) for 14-15 days. To derive A375 persister cells at high cell density, cells were seeded at 4,000/well in 96 well plates and after 24 hours were subjected to continuous treatment with 10 nM dabrafenib and 1 nM trametinib for 14-15 days. To derive BT474 low density persister cells, cells were seeded at 5,500 /well in 96 well plates and after 72 hours were subjected to continuous treatment with 2 μM lapatinib for 10 days. For all persister cell drug treatments, targeted therapy was refreshed every 3-4 days.

### Cell viability assays

Cell viability was evaluated by measuring ATP levels using CellTiter-Glo® Luminescent Cell Viability assay (CTG) (Promega, G7570) according to the manufacturer’s manual. Sample luminescence was measured in black optical-bottom plates (Corning, 39304) using the SpectraMax iD3 microplate reader and SoftMax Pro 7 Software.

### Single cell RNA seq analysis

To derive persister cells for the scRNAseq experiments, PC9 cells were adhered overnight and then treated with 100 nM erlotinib for 10 days. Parental PC9 cells were seeded on day 9 of persister cell treatment. The following day, or following persister cell derivation, 1 µM RSL3 was added to both persister and parental cells and samples were collected for single cell library preparation after 24 hours in RSL3. For panobinostat treatment, parental and persister cells were treated with 7.5 nM panobinostat for 48 hours prior to collection. Cells were analyzed with the 10x Genomics 3’ Chromium v3 platform for scRNAseq according to the manufacturer’s protocol. Libraries were generated and checked for quality and concentration using an Agilent Tapestation and Qubit respectively. Samples were combined and sequenced using an Illumina NovaSeq 6000 (Flow Cell Type: S4).

### ScRNAseq data mapping and processing

The Cell Ranger Single-Cell Software Suite (version 3.1.0) was used to align fastq files to the human reference genome “refdata-cellranger-GRCh38-3.0.0” with the “cellranger count” command. Mapped reads from individual samples were then merged as Seurat objects (Seurat version 3.1) (*60, 61*). We selected cells with greater than 1,000 and less than 7,500 features, and with less than 20% mitochondrial content for downstream analysis. Normalization and scaling were performed with the Seurat “SCTransform” command with cell cycle gene regression (*62, 63*). Downstream commands “RunPCA,” “RunUMAP,” “FindNeighbors,” and “FindClusters” were performed with default settings, with 30 dimensions used for “RunUMAP” and “FindNeighbors.” Differentially expressed genes were calculated with the Seurat “FindMarkers” command without expression or cell number thresholds. Gene set enrichment analysis was conducted with the ClusterProfiler R package (version 3.18.0) with default settings using Hallmarks and Oncogenic signatures gene sets (*64*). Pseudotime analysis was performed with Slingshot (version 2.14.0) with both persister cell clusters set as the starting clusters (*65*).

### Immunoblotting

Parental and persister cells were washed with PBS and lysed with RIPA buffer (ThermoFisher Scientific, 89900) with phosphatase inhibitor (ThermoFisher Scientific, 78420) and protease inhibitor (ThermoFisher Scientific, 78430). Lysates were centrifuged at 14,000 x *g* at 4 °C for 15 minutes, and protein concentration was determined using the Pierce BCA Protein Assay Kit (ThermoFisher Scientific, 23225). Lysates were combined with sample buffer (ThermoFisher Scientific, NP0007) and incubated at 70 °C for 10 minutes. Samples were run on SDS– PAGE gels (Bolt 4–12% Bis-Tris Gel, Life Technologies, NW04120BOX), with Chameleon Duo Pre-stained Protein Ladder (LICOR, #928-60000), and transferred to a nitrocellulose membrane using the iBLOT 2 Dry Blotting System (Life Technologies, IB21001). Membranes were blocked with 5% BSA (GeminiBio, 700-100P-1KG) for 1 hour at room temperature before overnight incubation at 4 °C. The following day, LICOR secondary antibodies were incubated with the membrane for 1 hour at room temperature, and membranes were imaged using the LICOR Odyssey Imaging System and Image Studio version 5.2. Loading controls were either β-Tubulin or vinculin as indicated. Antibody commercial sources were: β-Tubulin (Invitrogen, MA5-16308); Vinculin (Cell Signaling Technology, #4650); NRF2 (Cell Signaling Technology, #12721); KEAP1 (Cell Signaling Technology, #8047); SLC7A11 (Cell Signaling Technology, #12691); SLC3A2 (Cell Signaling Technology, #47213); GPX4 (Cell Signaling Technology, #59735); AIFM2/FSP1 (Cell Signaling Technology, #51676); DHODH (Cell Signaling Technology, #26381); VKORC1L1 (Cell Signaling Technology, #29458); IRDye 680RD Goat anti-Mouse IgG secondary antibody (LICOR, #926-68070); IRDye 800CW Goat anti-Rabbit IgG secondary antibody (LICOR, #926-32211).

### Glutathione measurement

Intracellular levels of total glutathione were measured using GSH-Glo Glutathione Assay (Promega, V6911) according to the Assay Procedure for Adherent Mammalian Cells in the manufacturer’s manual and normalized by protein levels measured by Pierce BCA Protein Assay Kit (ThermoFisher Scientific, 23225) after protein isolation from cells using RIPA Lysis and Extraction Buffer as previously described (ThermoFisher Scientific, 1338-43-8).

### Measurement of synergistic drug effects

Cells plated in 96-well plates were treated with 6 concentrations of each chemical. After treatment with the respective chemicals for 24 hours, cell viability was quantitated by CellTiter-Glo. Bliss synergy scores were calculated using SynergyFinder 3.0 (*66*).

### Intracellular ROS measurements

Intracellular ROS level of persister cells were quantitated by flow cytometry using CellROX® Deep Red Flow Cytometry Assay Kit (Life Technologies, C10491), according to the manufacturer’s manual. Briefly, persister cells were treated with 7.5 nM panobinostat or DMSO for 2 days under continuous erlotinib treatment and then subjected to flow cytometry. For a positive control, cells were treated with 200 μM tert-butyl hydroperoxide (TBHP) for 60 minutes prior to the flow cytometry assay. Trypsin-lifted cells were incubated with 1 μM CellROX Deep Red reagent for 60 minutes at 37 °C without light exposure. During the final 15 minutes of staining, 1 μM SYTOX® Blue Dead Cell stain (ThermoFisher Scientific, S34857) solution was added to cells, followed by immediate analysis by flow cytometry using 405 nm excitation for the SYTOX® Blue Dead Cell stain and 635 nm excitation for the CellROX Deep Red reagent. See supplementary figure S6 for gating strategy.

### Iron measurement

Cell Ferrous Iron Colorimetric Assay Kit (Elabscience, E-BC-K881-M) was used to detect intracellular labile ferrous iron (Fe^2+^) according to the manufacturer’s manual. PC9 parental and persister cells were treated with 7.5 nM panobinostat or DMSO control for 2 days. Cells were then trypsinized and counted to retrieve 5 million cells per replicate. Background measurement from a control solution was subtracted from each sample and iron concentrations were calculated from a standard curve.

### Statistical analyses

Statistical tests and graphing were performed with GraphPad Prism 9.3.1 except for synergy calculation which was performed by SynergyFinder 3.0 (*66*). Unless otherwise noted, P values were calculated using unpaired, two-tailed t-tests assuming unequal variance. For ferroptosis-driver and ferroptosis-suppressor gene set signature scoring comparisons, P values were calculated with the Mann-Whitney test.

## Acknowledgments

We thank the Flow Cytometry Core at the San Diego Center for AIDS Research (SD CFAR), an NIH-funded program (P30 AI036214), which is supported by the following NIH Institutes and Centers: NIAID, NCI, NHLBI, NIA, NICHD, NIDA, NIDCR, NIDDK, NIMH, NIMHD, NINR, FIC, and OAR. We also thank the San Diego Center for AIDS Research Flow Cytometry Core funded by National Institutes of Health P30 AI036214, VA San Diego Health Care System, and the San Diego Veterans Medical Research Foundation. This publication includes data generated at the UC San Diego Institute for Genomic Medicine Genomics Center utilizing an Illumina NovaSeq 6000 that was purchased with funding from a Nation Institutes of Health SIG grant (#S10 OD026929). BioRender.com was used to generate figure cartoons.

## Funding

National Institutes of Health training grant T32CA067754 (A.F.W.)

National Institutes of Health training grant T32CA009523 (A.F.W.)

National Institutes of Health/National Cancer Institute R01CA212767 (M.J.H.)

Department of Defense (DOD) Congressionally Directed Medical

Research Program (CDMRP) Melanoma Research Program (MRP) Idea Award HT9425-23-1-0719 (M.J.H.)

DOD CDMRP MRP Melanoma Academy Scholar Award HT9425-24-1-0288 (M.J.H.) Curebound Discovery Grant (M.J.H.)

American Cancer Society IRG #19-230-48 (M.J.H.)

UCSD Moores Cancer Center Specialized Cancer Center Support Grant National Institutes of Health/National Cancer Institute P30CA023100 (M.J.H.)

V Foundation for Cancer Research V Scholar Award V2021-035 (M.J.H.)

Bristol-Myers Squibb Melanoma Research Alliance Young Investigator Award (M.J.H.)

University of California Research Initiatives Cancer Research Coordinating Committee Seed Grant C23CR5537 (M.J.H.)

University of California Academic Senate Bridge Grant BG104446 (M.J.H.) Tower Cancer Research Foundation Career Development Grant (M.J.H.)

The Skin Cancer Foundation Ashley Trenner Research Grant Award (M.J.H.).

## Author contributions

Conceptualization: MH, AFW, AES, MJH

Methodology: MH, AFW, AES, AHN, DAG, CET

Investigation: MH, AFW, AES, AHN, DAG, CET, SC

Supervision: MJH

Writing—original draft: AFW, MH, AES, MJH

Writing—review & editing: AFW, AES, MJH

## Competing interests

MJH is a co-founder of Ferro Therapeutics, a subsidiary of BridgeBio Pharma, Inc.

## Data and materials availability

All data required to evaluate the conclusions in this paper are available in the main text or in the Supplementary Materials. The raw sequence data from this study are publicly available in the NCBI’s Gene Expression Omnibus (accession number: GSE303411).

## Supplementary Materials

This PDF file includes:

Figs. S1 to S6

Uncropped western blot images

Captions for Tables S1-S8

## References

M. Russo, M. Chen, E. Mariella, H. Peng, S. K. Rehman, E. Sancho, A. Sogari, T. S. Toh, N. Q. Balaban, E. Batlle, R. Bernards, M. J. Garnett, M. Hangauer, E. Leucci, J.-C. Marine, C. A. O’Brien, Y. Oren, E. E. Patton, C. Robert, S. M. Rosenberg, S. Shen, A. Bardelli, Cancer drug-tolerant persister cells: from biological questions to clinical opportunities. Nat. Rev. Cancer 24, 694–717 (2024).

H. Terai, S. Kitajima, D. S. Potter, Y. Matsui, L. G. Quiceno, T. Chen, T. Kim, M. Rusan, T. C. Thai, F. Piccioni, K. A. Donovan, N. Kwiatkowski, K. Hinohara, G. Wei, N. S. Gray, E. S. Fischer, K.-K. Wong, T. Shimamura, A. Letai, P. S. Hammerman, D. A. Barbie, ER Stress Signaling Promotes the Survival of Cancer “Persister Cells” Tolerant to EGFR Tyrosine Kinase Inhibitors. Cancer Res. 78, 1044–1057 (2018).

M. J. Hangauer, V. S. Viswanathan, M. J. Ryan, D. Bole, J. K. Eaton, A. Matov, J. Galeas, H. D. Dhruv, M. E. Berens, S. L. Schreiber, F. McCormick, M. T. McManus, Drug-tolerant persister cancer cells are vulnerable to GPX4 inhibition. Nature 551, 247–250 (2017).

V. S. Viswanathan, M. J. Ryan, H. D. Dhruv, S. Gill, O. M. Eichhoff, B. Seashore-Ludlow, S. D. Kaffenberger, J. K. Eaton, K. Shimada, A. J. Aguirre, S. R. Viswanathan, S. Chattopadhyay, P. Tamayo, W. S. Yang, M. G. Rees, S. Chen, Z. V. Boskovic, S. Javaid, C. Huang, X. Wu, Y.-Y. Tseng, E. M. Roider, D. Gao, J. M. Cleary, B. M. Wolpin, J. P. Mesirov, D. A. Haber, J. A. Engelman, J. S. Boehm, J. D. Kotz, C. S. Hon, Y. Chen, W. C. Hahn, M. P. Levesque, J. G. Doench, M. E. Berens, A. F. Shamji, P. A. Clemons, B. R. Stockwell, S. L. Schreiber, Dependency of a therapy-resistant state of cancer cells on a lipid peroxidase pathway. Nature 547, 453–457 (2017).

X. Sun, J. M. Bieber, H. Hammerlindl, R. J. Chalkley, K. H. Li, A. L. Burlingame, M. P. Jacobson, L. F. Wu, S. J. Altschuler, Modulating environmental signals to reveal mechanisms and vulnerabilities of cancer persisters. Sci. Adv. 8, eabi7711 (2022).

K. N. Shah, R. Bhatt, J. Rotow, J. Rohrberg, V. Olivas, V. E. Wang, G. Hemmati, M. M. Martins, A. Maynard, J. Kuhn, J. Galeas, H. J. Donnella, S. Kaushik, A. Ku, S. Dumont, G. Krings, H. J. Haringsma, L. Robillard, A. D. Simmons, T. C. Harding, F. McCormick, A. Goga, C. M. Blakely, T. G. Bivona, S. Bandyopadhyay, Aurora kinase A drives the evolution of resistance to third-generation EGFR inhibitors in lung cancer. Nat. Med. 25, 111–118 (2019).

Y. Oren, M. Tsabar, M. S. Cuoco, L. Amir-Zilberstein, H. F. Cabanos, J.-C. Hütter, B. Hu, P. I. Thakore, M. Tabaka, C. P. Fulco, W. Colgan, B. M. Cuevas, S. A. Hurvitz, D. J. Slamon, A. Deik, K. A. Pierce, C. Clish, A. N. Hata, E. Zaganjor, G. Lahav, K. Politi, J. S. Brugge, A. Regev, Cycling cancer persister cells arise from lineages with distinct programs. Nature 596, 576–582 (2021).

G. D. Guler, C. A. Tindell, R. Pitti, C. Wilson, K. Nichols, T. KaiWai Cheung, H.-J. Kim, M. Wongchenko, Y. Yan, B. Haley, T. Cuellar, J. Webster, N. Alag, G. Hegde, E. Jackson, T. L. Nance, P. G. Giresi, K.-B. Chen, J. Liu, S. Jhunjhunwala, J. Settleman, J.-P. Stephan, D. Arnott, M. Classon, Repression of Stress-Induced LINE-1 Expression Protects Cancer Cell Subpopulations from Lethal Drug Exposure. Cancer Cell 32, 221-237.e13 (2017).

S. Shen, S. Faouzi, A. Bastide, S. Martineau, H. Malka-Mahieu, Y. Fu, X. Sun, C. Mateus, E. Routier, S. Roy, L. Desaubry, F. André, A. Eggermont, A. David, J.-Y. Scoazec, S. Vagner, C. Robert, An epitranscriptomic mechanism underlies selective mRNA translation remodelling in melanoma persister cells. Nat. Commun. 10, 5713 (2019).

E. Dhimolea, R. de Matos Simoes, D. Kansara, A. Al’Khafaji, J. Bouyssou, X. Weng, S. Sharma, J. Raja, P. Awate, R. Shirasaki, H. Tang, B. J. Glassner, Z. Liu, D. Gao, J. Bryan, S. Bender, J. Roth, M. Scheffer, R. Jeselsohn, N. S. Gray, I. Georgakoudi, F. Vazquez, A. Tsherniak, Y. Chen, A. Welm, C. Duy, A. Melnick, B. Bartholdy, M. Brown, A. C. Culhane, C. S. Mitsiades, An Embryonic Diapause-like Adaptation with Suppressed Myc Activity Enables Tumor Treatment Persistence. Cancer Cell 39, 240-256.e11 (2021).

M. Vinogradova, V. S. Gehling, A. Gustafson, S. Arora, C. A. Tindell, C. Wilson, K. E. Williamson, G. D. Guler, P. Gangurde, W. Manieri, J. Busby, E. M. Flynn, F. Lan, H. Kim, S. Odate, A. G. Cochran, Y. Liu, M. Wongchenko, Y. Yang, T. K. Cheung, T. M. Maile, T. Lau, M. Costa, G. V. Hegde, E. Jackson, R. Pitti, D. Arnott, C. Bailey, S. Bellon, R. T. Cummings, B. K. Albrecht, J.-C. Harmange, J. R. Kiefer, P. Trojer, M. Classon, An inhibitor of KDM5 demethylases reduces survival of drug-tolerant cancer cells. Nat. Chem. Biol. 12, 531–538 (2016).

W.-H. Yang, C.-K. C. Ding, T. Sun, G. Rupprecht, C.-C. Lin, D. Hsu, J.-T. Chi, The Hippo Pathway Effector TAZ Regulates Ferroptosis in Renal Cell Carcinoma. Cell Rep. 28, 2501-2508.e4 (2019).

J. Wu, A. M. Minikes, M. Gao, H. Bian, Y. Li, B. R. Stockwell, Z.-N. Chen, X. Jiang, Intercellular interaction dictates cancer cell ferroptosis via NF2–YAP signalling. Nature 572, 402–406 (2019).

W.-H. Yang, Z. Huang, J. Wu, C.-K. C. Ding, S. K. Murphy, J.-T. Chi, A TAZ–ANGPTL4–NOX2 Axis Regulates Ferroptotic Cell Death and Chemoresistance in Epithelial Ovarian Cancer. Mol. Cancer Res. 18, 79–90 (2020).

S. J. Dixon, J. A. Olzmann, The cell biology of ferroptosis. Nat. Rev. Mol. Cell Biol. 25, 424–442 (2024).

S. Müller, F. Sindikubwabo, T. Cañeque, A. Lafon, A. Versini, B. Lombard, D. Loew, T.-D. Wu, C. Ginestier, E. Charafe-Jauffret, A. Durand, C. Vallot, S. Baulande, N. Servant, R. Rodriguez, CD44 regulates epigenetic plasticity by mediating iron endocytosis. Nat. Chem. 12, 929–938 (2020).

A. Schwab, Z. Rao, J. Zhang, A. Gollowitzer, K. Siebenkäs, N. Bindel, E. D’Avanzo, R. van Roey, Y. Hajjaj, E. Özel, I. Armstark, L. Bereuter, F. Su, J. Grander, E. Bonyadi Rad, A. Groenewoud, F. B. Engel, G. W. Bell, W. S. Henry, J. P. F. Angeli, M. P. Stemmler, S. Brabletz, A. Koeberle, T. Brabletz, Zeb1 mediates EMT/plasticity-associated ferroptosis sensitivity in cancer cells by regulating lipogenic enzyme expression and phospholipid composition. Nat. Cell Biol. 26, 1470–1481 (2024).

Y. Wang, M. Hu, J. Cao, F. Wang, J. R. Han, T. W. Wu, L. Li, J. Yu, Y. Fan, G. Xie, H. Lian, Y. Cao, N. Naowarojna, X. Wang, Y. Zou, ACSL4 and polyunsaturated lipids support metastatic extravasation and colonization. Cell 188, 412-429.e27 (2025).

R. Rodriguez, S. L. Schreiber, M. Conrad, Persister cancer cells: Iron addiction and vulnerability to ferroptosis. Mol. Cell 82, 728–740 (2022).

M.-J. Nokin, E. Darbo, E. Richard, S. San José, S. de Hita, V. Prouzet-Mauleon, B. Turcq, L. Gerardelli, R. Crake, V. Velasco, B. Koopmansch, F. Lambert, J. Y. Xue, B. Sang, J. Horne, E. Ziemons, A. Villanueva, Blomme, M. Herfs, D. Cataldo, O. Calvayrac, P. Porporato, E. Nadal, P. Lito, P. A. Jänne, B. Ricciuti, M. M. Awad, C. Ambrogio, D. Santamaría, In vivo vulnerabilities to GPX4 and HDAC inhibitors in drug-persistent versus drug-resistant BRAFV600E lung adenocarcinoma. Cell Rep. Med. 5, 101663 (2024).

T. Cañeque, L. Baron, S. Müller, A. Carmona, L. Colombeau, A. Versini, S. Solier, C. Gaillet, F. Sindikubwabo, J. L. Sampaio, M. Sabatier, E. Mishima, A. Picard-Bernes, L. Syx, N. Servant, B. Lombard, D. Loew, J. Zheng, B. Proneth, L. K. Thoidingjam, L. Grimaud, C. S. Fraser, K. J. Szylo, E. Der Kazarian, C. Bonnet, E. Charafe-Jauffret, C. Ginestier, P. Santofimia-Castaño, M. Estaras, N. Dusetti, J. L. Iovanna, A. S. Cunha, G. Pittau, P. Hammel, D. Tzanis, S. Bonvalot, S. Watson, V. Gandon, A. Upadhyay, D. A. Pratt, F. P. Freitas, J. P. Friedmann Angeli, B. R. Stockwell, M. Conrad, J. M. Ubellacker, R. Rodriguez, Activation of lysosomal iron triggers ferroptosis in cancer. Nature 642, 492–500 (2025).

H. Konishi, Y. Haga, Y. Lin, H. Tsujino, K. Higashisaka, Y. Tsutsumi, Osimertinib-tolerant lung cancer cells are susceptible to ferroptosis. Biochem. Biophys. Res. Commun. 641, 116–122 (2023).

G. V. Echeverria, Z. Ge, S. Seth, X. Zhang, S. Jeter-Jones, X. Zhou, S. Cai, Y. Tu, A. McCoy, M. Peoples, Y. Sun, H. Qiu, Q. Chang, C. Bristow, A. Carugo, J. Shao, X. Ma, A. Harris, P. Mundi, R. Lau, V. Ramamoorthy, Y. Wu, M. J. Alvarez, A. Califano, S. L. Moulder, W. F. Symmans, J. R. Marszalek, T. P. Heffernan, J. T. Chang, H. Piwnica-Worms, Resistance to neoadjuvant chemotherapy in triple-negative breast cancer mediated by a reversible drug-tolerant state. Sci. Transl. Med. 11, eaav0936 (2019).

S. Tau, M. D. Chamberlin, H. Yang, J. D. Marotti, P. C. Muskus, A. M. Roberts, M. M. Carmichael, L. Cressey, C. P. C. Dragnev, E. Demidenko, R. A. Hampsch, S. M. Soucy, F. W. Kolling, K. S. Samkoe, J. V. Alvarez, A. N. Kettenbach, T. W. Miller, Oxidative Phosphorylation Is a Metabolic Vulnerability of Endocrine Therapy–Tolerant Persister Cells in ER+ Breast Cancer. Cancer Res. 85, 1145–1161 (2025).

S. Shen, S. Faouzi, S. Souquere, S. Roy, E. Routier, C. Libenciuc, F. André, G. Pierron, J.-Y. Scoazec, C. Robert, Melanoma Persister Cells Are Tolerant to BRAF/MEK Inhibitors via ACOX1-Mediated Fatty Acid Oxidation. Cell Rep. 33 (2020).

W. Liu, B. Chakraborty, R. Safi, D. Kazmin, C. Chang, D. P. McDonnell, Dysregulated cholesterol homeostasis results in resistance to ferroptosis increasing tumorigenicity and metastasis in cancer. Nat. Commun. 12, 5103 (2021).

J. Yi, J. Zhu, J. Wu, C. B. Thompson, X. Jiang, Oncogenic activation of PI3K-AKT-mTOR signaling suppresses ferroptosis via SREBP-mediated lipogenesis. Proc. Natl. Acad. Sci. 117, 31189–31197 (2020).

J. Y. Cao, S. J. Dixon, Mechanisms of ferroptosis. Cell. Mol. Life Sci. 73, 2195–2209 (2016).

J. Wu, Z. Feng, L. Chen, Y. Li, H. Bian, J. Geng, Z.-H. Zheng, X. Fu, Z. Pei, Y. Qin, L. Yang, Y. Zhao, K. Wang, R. Chen, Q. He, G. Nan, X. Jiang, Z.-N. Chen, P. Zhu, TNF antagonist sensitizes synovial fibroblasts to ferroptotic cell death in collagen-induced arthritis mouse models. Nat. Commun. 13, 676 (2022).

A. Tarangelo, L. Magtanong, K. T. Bieging-Rolett, Y. Li, J. Ye, L. D. Attardi, S. J. Dixon, p53 Suppresses Metabolic Stress-Induced Ferroptosis in Cancer Cells. Cell Rep. 22, 569–575 (2018).

N. Zhou, X. Yuan, Q. Du, Z. Zhang, X. Shi, J. Bao, Y. Ning, L. Peng, FerrDb V2: update of the manually curated database of ferroptosis regulators and ferroptosis-disease associations. Nucleic Acids Res. 51, D571–D582 (2023).

T. M. Seibt, B. Proneth, M. Conrad, Role of GPX4 in ferroptosis and its pharmacological implication. Free Radic. Biol. Med. 133, 144–152 (2019).

S. Doll, F. P. Freitas, R. Shah, M. Aldrovandi, M. C. da Silva, I. Ingold, A. Goya Grocin, T. N. Xavier da Silva, E. Panzilius, C. H. Scheel, A. Mourão, K. Buday, M. Sato, J. Wanninger, T. Vignane, V. Mohana, M. Rehberg, A. Flatley, A. Schepers, A. Kurz, D. White, M. Sauer, M. Sattler, E. W. Tate, W. Schmitz, A. Schulze, V. O’Donnell, B. Proneth, G. M. Popowicz, D. A. Pratt, J. P. F. Angeli, M. Conrad, FSP1 is a glutathione-independent ferroptosis suppressor. Nature 575, 693–698 (2019).

K. Bersuker, J. M. Hendricks, Z. Li, L. Magtanong, B. Ford, P. H. Tang, M. A. Roberts, B. Tong, T. J. Maimone, R. Zoncu, M. C. Bassik, D. K. Nomura, S. J. Dixon, J. A. Olzmann, The CoQ oxidoreductase FSP1 acts parallel to GPX4 to inhibit ferroptosis. Nature 575, 688–692 (2019).

C. Mao, X. Liu, Y. Zhang, G. Lei, Y. Yan, H. Lee, P. Koppula, S. Wu, L. Zhuang, B. Fang, M. V. Poyurovsky, K. Olszewski, B. Gan, DHODH-mediated ferroptosis defence is a targetable vulnerability in cancer. Nature 593, 586–590 (2021).

X. Yang, Z. Wang, F. Zandkarimi, Y. Liu, S. Duan, Z. Li, N. Kon, Z. Zhang, X. Jiang, B. R. Stockwell, W. Gu, Regulation of VKORC1L1 is critical for p53-mediated tumor suppression through vitamin K metabolism. Cell Metab. 35, 1474-1490.e8 (2023).

R. Bian, Y. Shang, N. Xu, B. Liu, Y. Ma, H. Li, J. Chen, Q. Yao, HDAC inhibitor enhances ferroptosis susceptibility of AML cells by stimulating iron metabolism. Cell. Signal. 127, 111583 (2025).

Z. Yang, W. Su, Q. Zhang, L. Niu, B. Feng, Y. Zhang, F. Huang, J. He, Q. Zhou, X. Zhou, L. Ma, J. Zhou, Y. Wang, W. Xiong, J. Xiang, Z. Hu, Q. Zhan, B. Yao, Lactylation of HDAC1 Confers Resistance to Ferroptosis in Colorectal Cancer. Adv. Sci. 12, 2408845 (2025).

H. Yang, L. Zhao, Y. Gao, F. Yao, T. M. Marti, R. A. Schmid, R.-W. Peng, Pharmacotranscriptomic Analysis Reveals Novel Drugs and Gene Networks Regulating Ferroptosis in Cancer. Cancers 12, 3273 (2020).

M. Zille, A. Kumar, N. Kundu, M. W. Bourassa, V. S. C. Wong, D. Willis, S. S. Karuppagounder, R. R. Ratan, Ferroptosis in Neurons and Cancer Cells Is Similar But Differentially Regulated by Histone Deacetylase Inhibitors. eNeuro 6 (2019).

K. Miyamoto, M. Watanabe, S. Boku, M. Sukeno, M. Morita, H. Kondo, K. Sakaguchi, T. Taguchi, T. Sakai, xCT Inhibition Increases Sensitivity to Vorinostat in a ROS-Dependent Manner. Cancers 12, 827 (2020).

T. Alothaim, M. Charbonneau, X. Tang, HDAC6 inhibitors sensitize non-mesenchymal triple-negative breast cancer cells to cysteine deprivation. Sci. Rep. 11, 10956 (2021).

T. Zhang, B. Sun, C. Zhong, K. Xu, Z. Wang, P. Hofman, T. Nagano, A. Legras, D. Breadner, B. Ricciuti, D. Divisi, R. A. Schmid, R.-W. Peng, H. Yang, F. Yao, Targeting histone deacetylase enhances the therapeutic effect of Erastin-induced ferroptosis in EGFR-activating mutant lung adenocarcinoma. Transl. Lung Cancer Res. 10 (2021).

T. Oliveira, E. Hermann, D. Lin, W. Chowanadisai, E. Hull, M. Montgomery, HDAC inhibition induces EMT and alterations in cellular iron homeostasis to augment ferroptosis sensitivity in SW13 cells. Redox Biol. 47, 102149 (2021).

M. Gao, J. Deng, F. Liu, A. Fan, Y. Wang, H. Wu, D. Ding, D. Kong, Z. Wang, D. Peer, Y. Zhao, Triggered ferroptotic polymer micelles for reversing multidrug resistance to chemotherapy. Biomaterials 223, 119486 (2019).

T. Ishida, T. Takahashi, Y. Kurokawa, T. Nishida, S. Hirota, S. Serada, M. Fujimoto, T. Naka, R. Teranishi, T. Saito, K. Yamashita, K. Tanaka, K. Yamamoto, T. Makino, M. Yamasaki, K. Nakajima, H. Eguchi, Y. Doki, Targeted therapy for drug-tolerant persister cells after imatinib treatment for gastrointestinal stromal tumours. Br. J. Cancer 125, 1511–1522 (2021).

J. H. You, J. Lee, J.-L. Roh, Mitochondrial pyruvate carrier 1 regulates ferroptosis in drug-tolerant persister head and neck cancer cells via epithelial-mesenchymal transition. Cancer Lett. 507, 40–54 (2021).

H. Kalkavan, M. J. Chen, J. C. Crawford, G. Quarato, P. Fitzgerald, S. W. G. Tait, C. R. Goding, D. R. Green, Sublethal cytochrome c release generates drug-tolerant persister cells. Cell 185, 3356-3374.e22 (2022).

X. Zhang, Y. Ma, J. Ma, L. Yang, Q. Song, H. Wang, G. Lv, Glutathione Peroxidase 4 as a Therapeutic Target for Anti-Colorectal Cancer Drug-Tolerant Persister Cells. Front. Oncol. 12 (2022).

D. Nolfi-Donegan, A. Braganza, S. Shiva, Mitochondrial electron transport chain: Oxidative phosphorylation, oxidant production, and methods of measurement. Redox Biol. 37, 101674 (2020).

M.-Q. Shi, Y. Xu, X. Fu, D.-S. Pan, X.-P. Lu, Y. Xiao, Y.-Z. Jiang, Advances in targeting histone deacetylase for treatment of solid tumors. J. Hematol. Oncol.J Hematol Oncol 17, 37 (2024).

R. Jenke, D. Oliinyk, T. Zenz, J. Körfer, L. Schäker-Hübner, F. K. Hansen, F. Lordick, F. Meier-Rosar, A. Aigner, T. Büch, HDAC inhibitors activate lipid peroxidation and ferroptosis in gastric cancer. Biochem. Pharmacol. 225, 116257 (2024).

E. Karaj, S. H. Sindi, N. Kuganesan, R. A. Koranne, J. R. Knoff, A. W. James, Y. Fu, L. N. Kotsull, M. K. Pflum, Z. Shah, W. R. Taylor, L. M. V. Tillekeratne, First-in-Class Dual Mechanism Ferroptosis-HDAC Inhibitor Hybrids. J. Med. Chem. 65, 14764–14791 (2022).

S. Ramos, E. Hartenian, J. C. Santos, P. Walch, P. Broz, NINJ1 induces plasma membrane rupture and release of damage-associated molecular pattern molecules during ferroptosis. EMBO J. 43, 1164–1186 (2024).

L. Pedrera, R. A. Espiritu, U. Ros, J. Weber, A. Schmitt, J. Stroh, S. Hailfinger, S. von Karstedt, A.J. García-Sáez, Ferroptotic pores induce Ca2+ fluxes and ESCRT-III activation to modulate cell death kinetics. Cell Death Differ. 28, 1644–1657 (2021).

E. Dai, L. Meng, R. Kang, X. Wang, D. Tang, ESCRT-III–dependent membrane repair blocks ferroptosis. Biochem. Biophys. Res. Commun. 522, 415–421 (2020).

L. J. Yant, Q. Ran, L. Rao, H. Van Remmen, T. Shibatani, J. G. Belter, L. Motta, A. Richardson, T. A. Prolla, The selenoprotein GPX4 is essential for mouse development and protects from radiation and oxidative damage insults. Free Radic. Biol. Med. 34, 496–502 (2003).

H. Imai, F. Hirao, T. Sakamoto, K. Sekine, Y. Mizukura, M. Saito, T. Kitamoto, M. Hayasaka, K. Hanaoka, Y. Nakagawa, Early embryonic lethality caused by targeted disruption of the mouse PHGPx gene. Biochem. Biophys. Res. Commun. 305, 278–286 (2003).

S.-E. Yoo, L. Chen, R. Na, Y. Liu, C. Rios, H. Van Remmen, A. Richardson, Q. Ran, Gpx4 ablation in adult mice results in a lethal phenotype accompanied by neuronal loss in brain. Free Radic. Biol. Med. 52, 1820–1827 (2012).

T. Stuart, A. Butler, P. Hoffman, C. Hafemeister, E. Papalexi, W. M. Mauck, Y. Hao, M. Stoeckius, P. Smibert, R. Satija, Comprehensive Integration of Single-Cell Data. Cell 177, 1888-1902.e21 (2019).

A. Butler, P. Hoffman, P. Smibert, E. Papalexi, R. Satija, Integrating single-cell transcriptomic data across different conditions, technologies, and species. Nat. Biotechnol. 36, 411–420 (2018).

C. Hafemeister, R. Satija, Normalization and variance stabilization of single-cell RNA-seq data using regularized negative binomial regression. Genome Biol. 20, 296 (2019).

M. S. Kowalczyk, I. Tirosh, D. Heckl, T. N. Rao, A. Dixit, B. J. Haas, R. K. Schneider, A. J. Wagers, B. L. Ebert, A. Regev, Single-cell RNA-seq reveals changes in cell cycle and differentiation programs upon aging of hematopoietic stem cells. Genome Res. 25, 1860–1872 (2015).

G. Yu, L.-G. Wang, Y. Han, Q.-Y. He, clusterProfiler: an R Package for Comparing Biological Themes Among Gene Clusters. OMICS J. Integr. Biol. 16, 284–287 (2012).

K. Street, D. Risso, R. B. Fletcher, D. Das, J. Ngai, N. Yosef, E. Purdom, S. Dudoit, Slingshot: cell lineage and pseudotime inference for single-cell transcriptomics. BMC Genomics 19, 477 (2018).

A. Ianevski, A. K. Giri, T. Aittokallio, SynergyFinder 3.0: an interactive analysis and consensus interpretation of multi-drug synergies across multiple samples. Nucleic Acids Res. 50, W739–W743 (2022).

