## Supplementary materials for "FSP1 and histone deacetylases suppress cancer persister cell ferroptosis"

### Cancer persister cell ferroptosis sensitivity is driven by oxidative stress

Masayoshi Higuchi *et al.*

**This PDF file includes:**

Figs. S1 to S6
Uncropped western blot images
Captions for Supplementary tables S1-S8

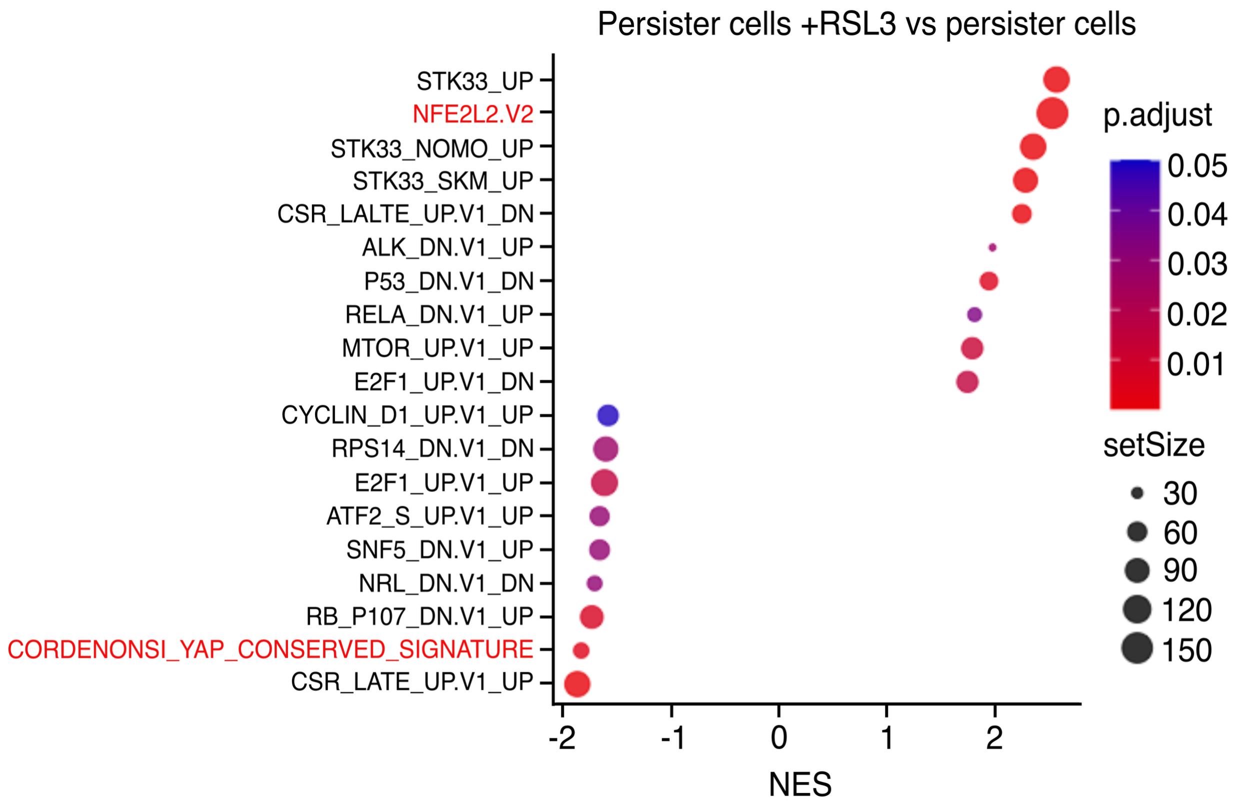


**Fig. S1. Enriched Oncogenic Signature gene sets between persister cells treated with or without RSL3.** Positive Normalized Enrichment Score (NES) values indicate increased gene set expression in cells treated with RSL3. NFE2L2.V2 refers to the NRF2 gene set.

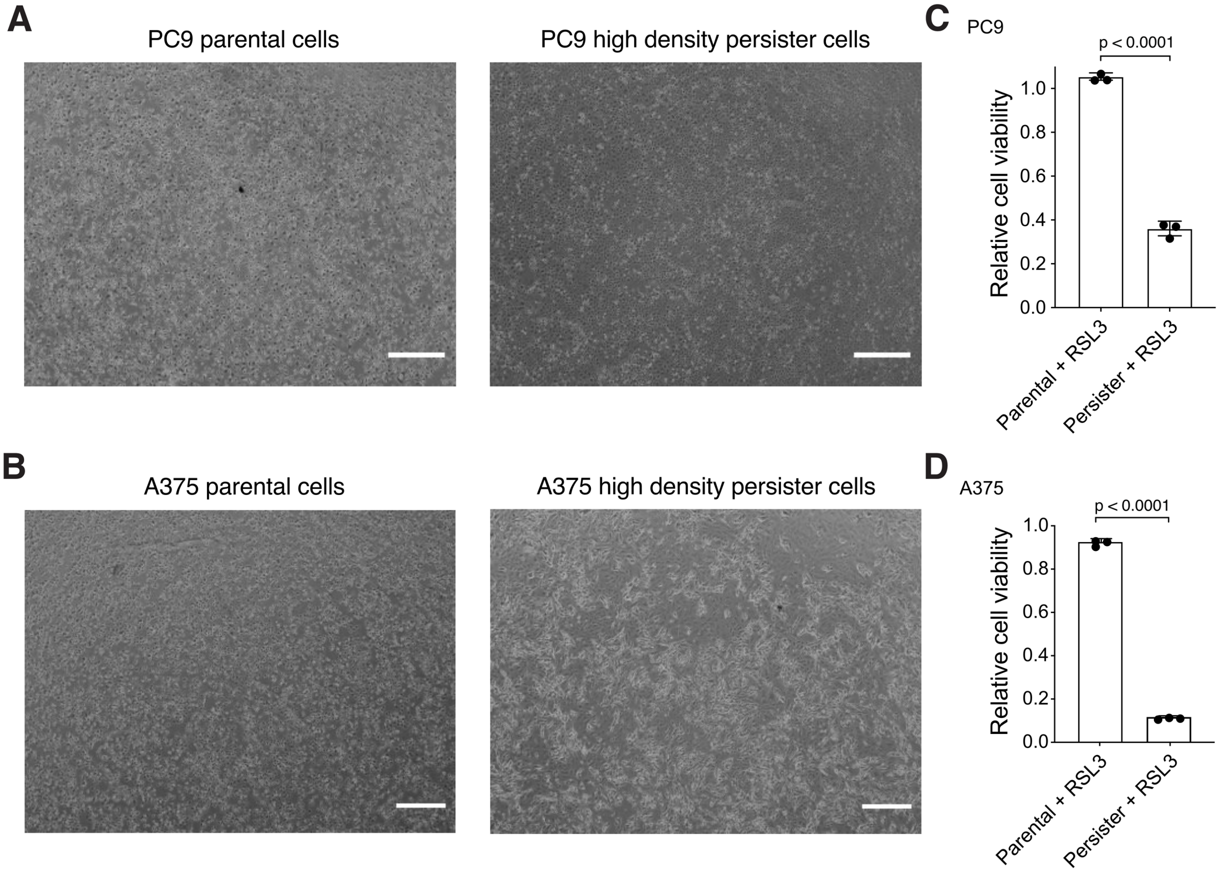


**Fig. S2. Persister cells remain sensitized to ferroptosis at high confluency.** (**A** and **B**) High density persister cells derived from 70 nM erlotinib for PC9 cells and 10 nM dabrafenib and 1 nM trametinib for A375 cells. Parental cells of similar density are also shown. See Materials and Methods. (**C** and **D**) Cell viability of PC9 and A375 high density persister cells and density-matched parental cells treated with 1 μM RSL3 for 24 hours. *n* = 3 biological replicates; mean ± s.d. is shown; P values calculated with two-tailed Student's t-test. Scale bar is 100 μm.

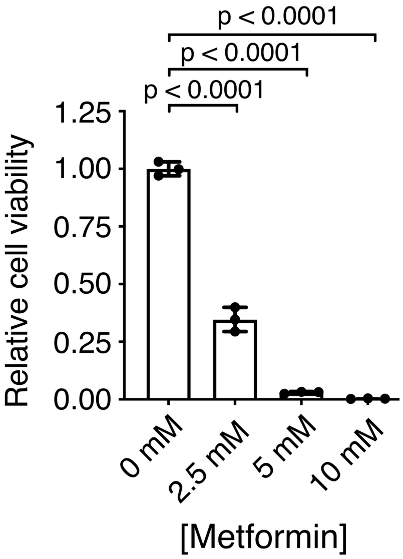


**Fig. S3. Metformin is toxic to persister cells.** PC9 persister cells derived from treatment with 70 nM erlotinib for 10 days were cotreated with the indicated metformin concentrations throughout drug treatment. $n$ = 3 biological replicates; mean ± s.d. is shown; P values calculated with two-tailed Student's t-test.

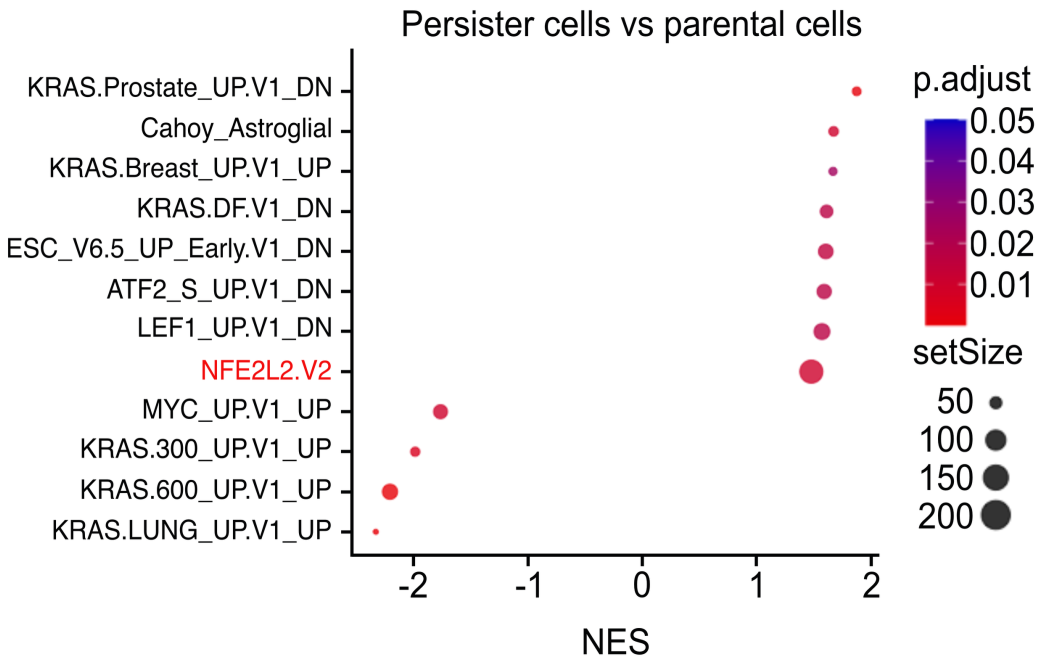


**Fig. S4. Enriched Oncogenic Signatures gene sets between PC9 parental and persister cells.** Positive Normalized Enrichment Score (NES) values indicate gene sets upregulated in persister cells. NFE2L2.V2 refers to the NRF2 gene set.

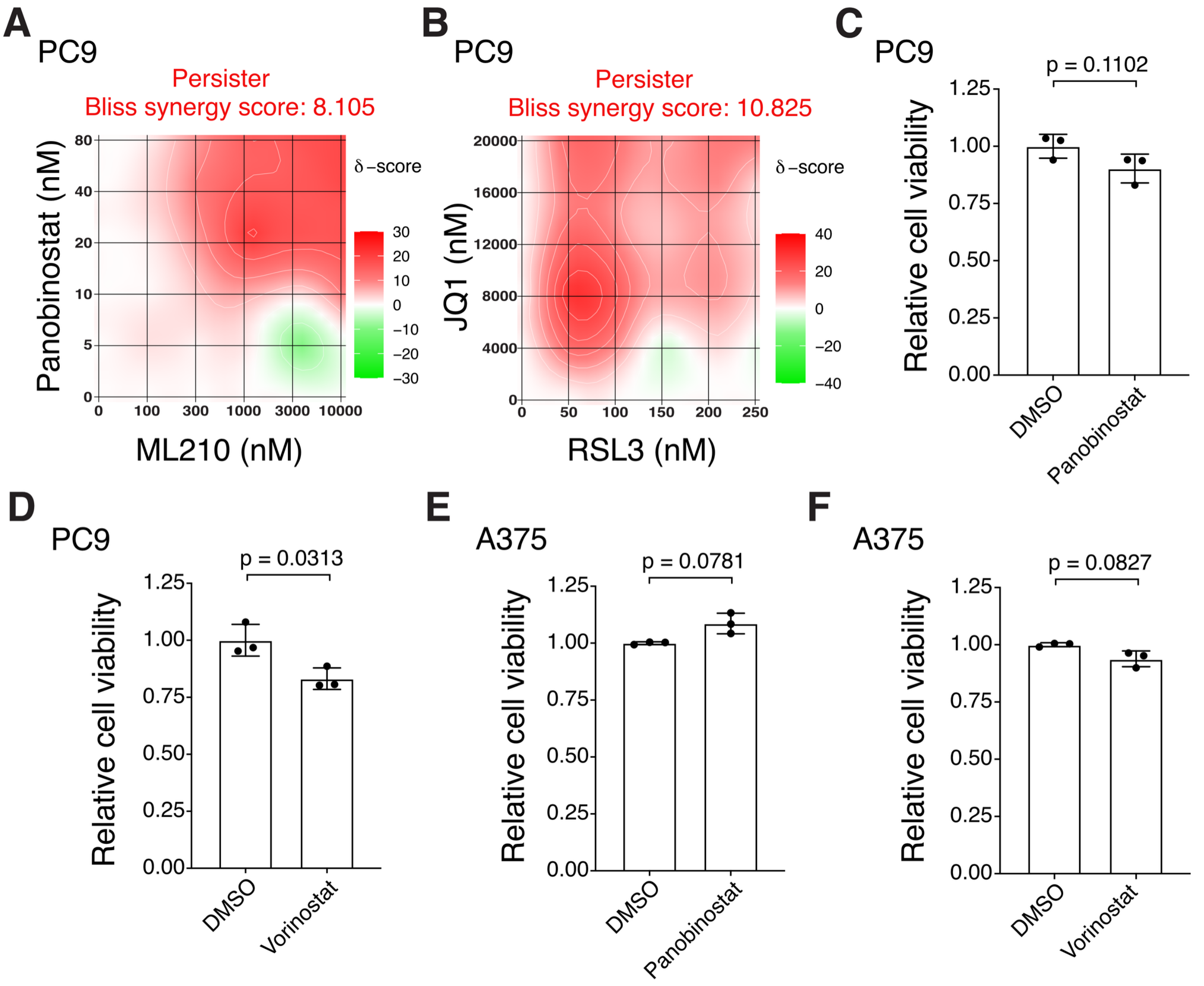


**Fig. S5. Additional data demonstrating HDAC and BRD4 inhibitors synergize with GPX4 inhibitors to kill persister cells.** (**A** and **B**) Heatmaps of synergy between GPX4 inhibitors ML210 (**A**) or RSL3 (**B**) and HDAC inhibitor panobinostat (**A**) or BRD4 inhibitor JQ1 (**B**) following 24-hour cotreatment of pre-derived PC9 persister cells. Bliss synergy score calculated with SynergyFinder 3.0. (**C** to **F**) Persister cells are insensitive to HDAC inhibitor concentrations which sensitize persister cells to ferroptosis. PC9 and A375 persister cell viability was measured following 48 hour treatment with 7.5 nM panobinostat (**C**), 100 nM vorinostat (**D**), 5 nM panobinostat (**E**), or 1 µM vorinostat (**F**). (**C-F**) $n$ = 3 biological replicates; mean ± s.d. is shown; P values calculated with two-tailed Student's t-test.

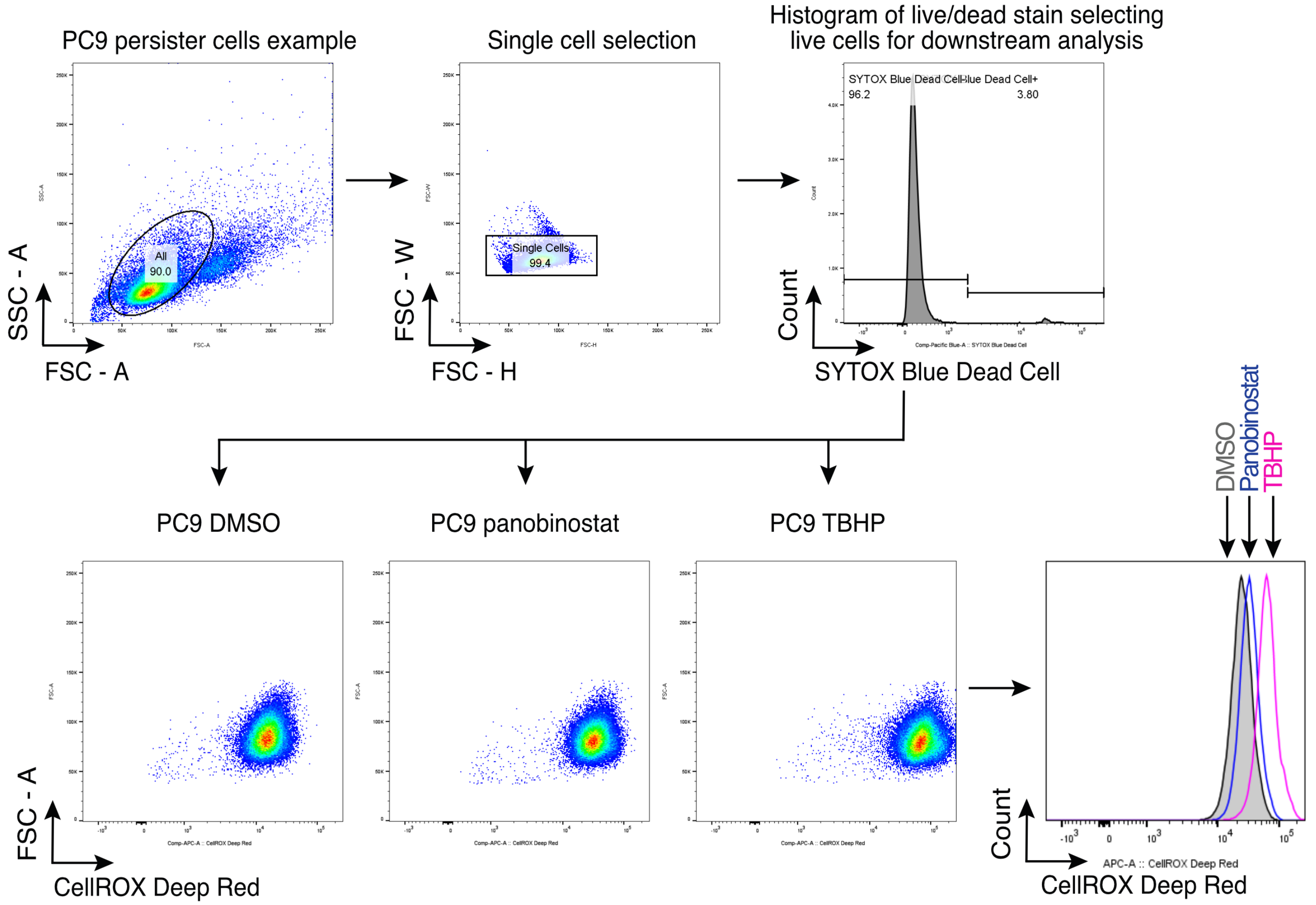


**Fig. S6. Flow cytometry schematic for CellROX Deep Red oxidative stress assay.** Gating strategy used for Fig. 4G, where geometric means of CellROX deep red signal from live, single cells are graphed. PC9 persister cells were co-treated with DMSO or 7.5 nM panobinostat for 48 hours. Positive control cells were treated with 200 μM tert-butyl hydroperoxide (TBHP) for 1 hour.

**Uncropped western blot images**

For uncropped western blot images, red boxes indicate bands shown in the main figure while blue boxes indicate loading controls that were not included in the main figures.

**Fig. 2A**

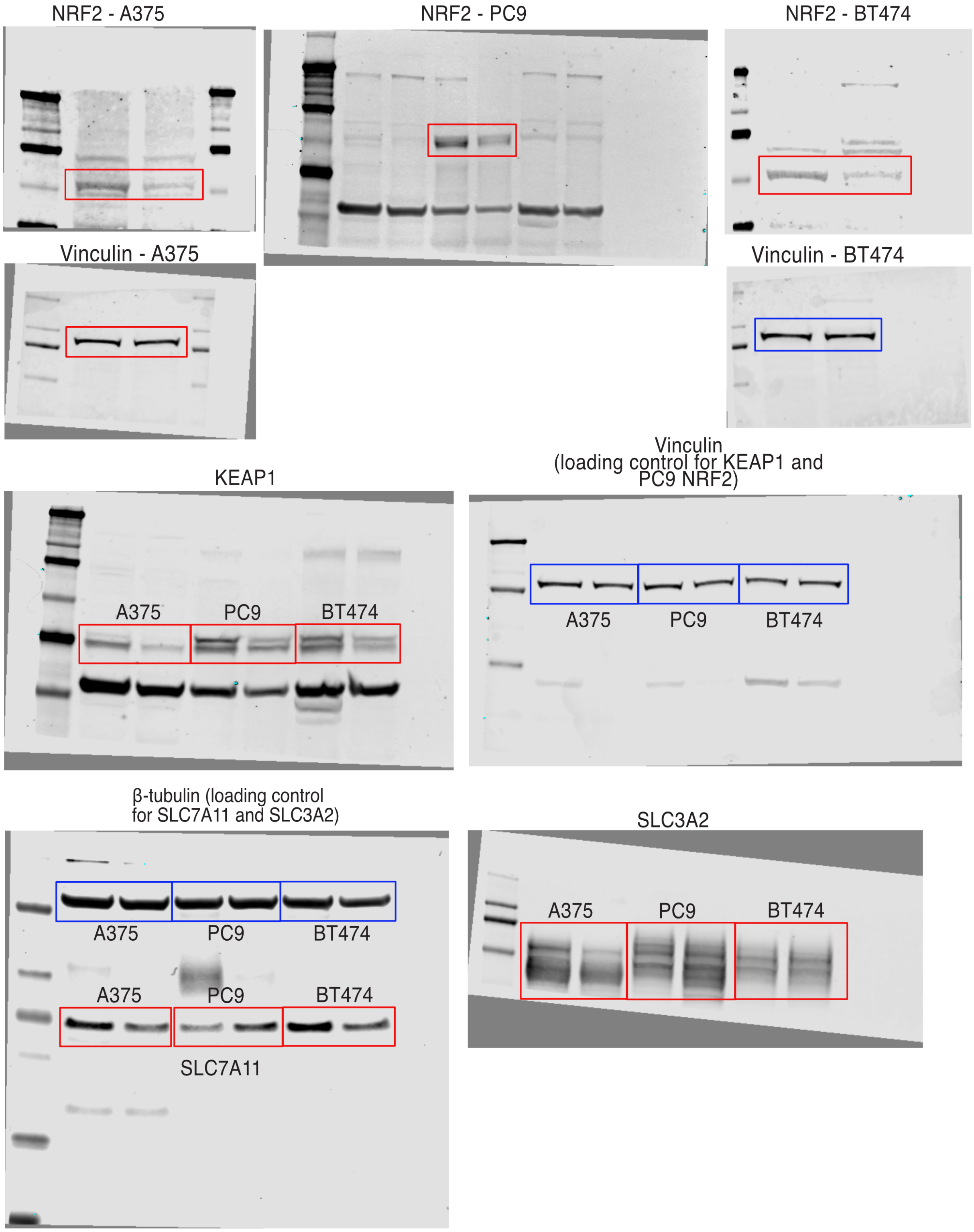

**Fig. 2D**

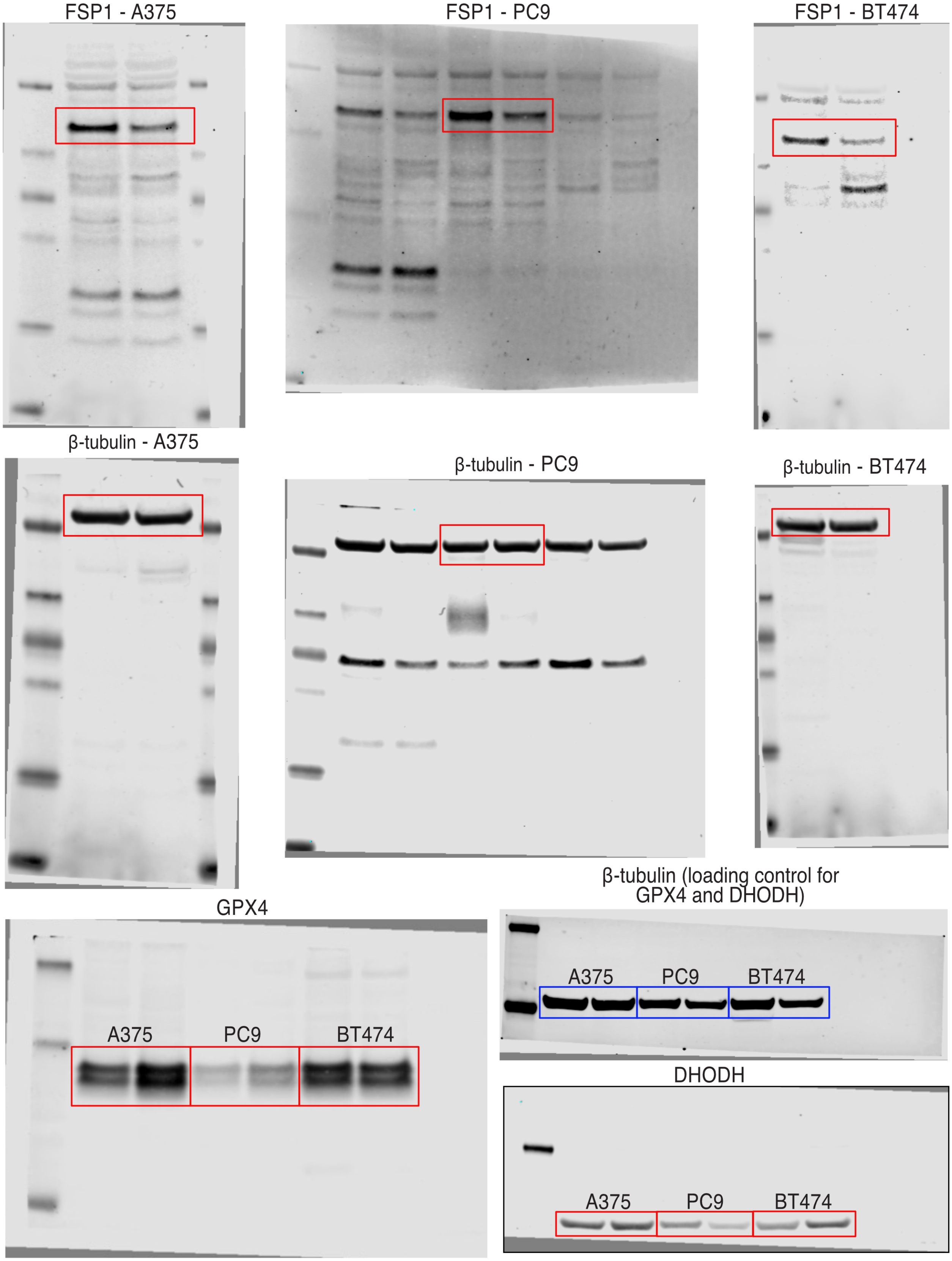
FSP1 - A375
FSP1 - PC9
FSP1 - BT474
β-tubulin - A375
β-tubulin - PC9
β-tubulin - BT474
β-tubulin (loading control for GPX4 and DHODH)
A375
PC9
BT474
GPX4
A375
PC9
BT474
DHODH
A375
PC9
BT474

**Fig. 2D (continued)**

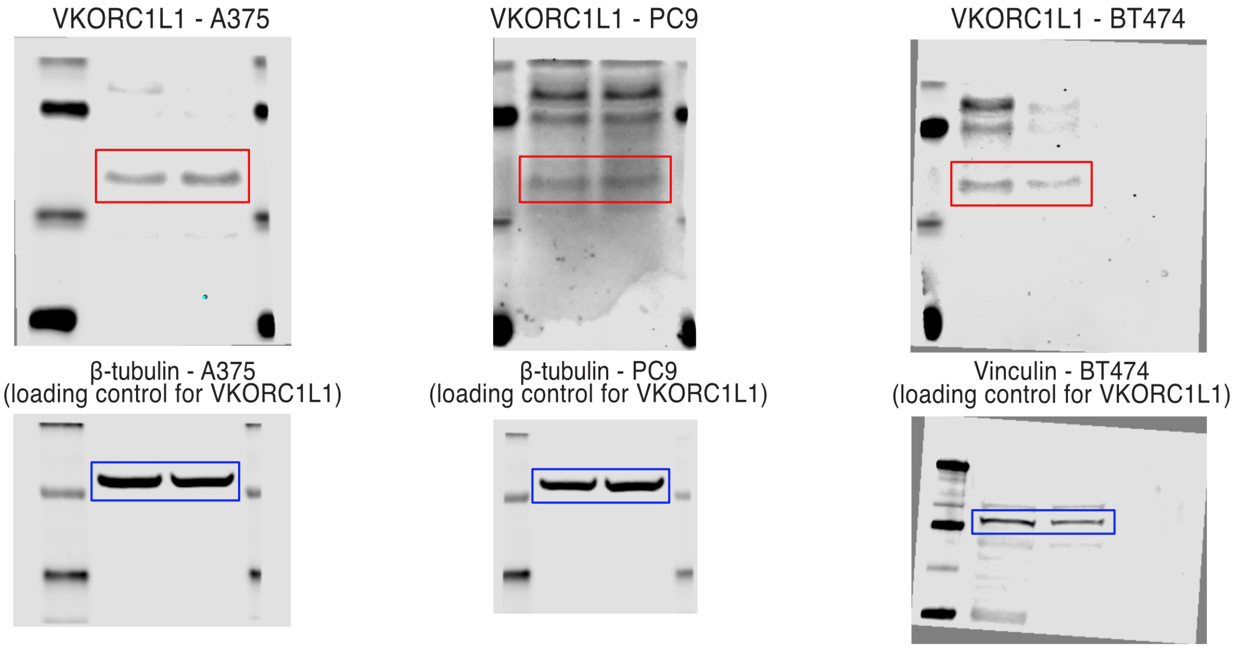


### Captions for Tables S1-S8

**Table S1.** Differentially expressed genes between PC9 persister cells treated with RSL3 versus without RSL3.

**Table S2.** Enriched Hallmarks and Oncogenic Signatures gene sets between PC9 persister cells treated with RSL3 versus without RSL3.

**Table S3.** Differentially expressed genes between PC9 persister cell clusters treated with RSL3 versus without RSL3.

**Table S4.** Enriched Hallmarks and Oncogenic Signatures gene sets between PC9 persister cell clusters treated with RSL3 versus without RSL3.

**Table S5.** Differentially expressed genes between PC9 persister and parental cells.

**Table S6.** Enriched Hallmarks and Oncogenic Signatures gene sets between PC9 persister and parental cells.

**Table S7.** Differentially expressed genes between PC9 persister cells treated with panobinostat versus without panobinostat.

**Table S8.** Enriched Hallmarks gene sets between PC9 persister cells treated with panobinostat versus without panobinostat.
